# Hypergraph factorisation for multi-tissue gene expression imputation

**DOI:** 10.1101/2022.07.31.502211

**Authors:** Ramon Viñas, Chaitanya K. Joshi, Dobrik Georgiev, Bianca Dumitrascu, Eric R. Gamazon, Pietro Liò

## Abstract

Integrating gene expression across scales and tissues is crucial for understanding the biological mechanisms that drive disease and characterise homeostasis. However, traditional multi-tissue integration methods cannot handle uncollected tissues or rely on genotype information, which is subject to privacy concerns and often unavailable. To address these challenges, we present HYFA (**Hy**pergraph **Fa**ctorisation), a novel method for joint imputation of multi-tissue and cell-type gene expression. HYFA imputes tissue-specific gene expression via a specialised graph neural network operating on a hypergraph of individuals, metagenes, and tissues. HYFA is genotype-agnostic, supports a variable number of collected tissues per individual, and imposes strong inductive biases to leverage the shared regulatory architecture of tissues. In performance comparison on data from the Genotype Tissue Expression project, HYFA achieves superior performance over existing transcriptome imputation methods, especially when multiple reference tissues are available. Through transfer learning on a paired single-nucleus RNA-seq (snRNA-seq) dataset, we further show that HYFA can accurately resolve cell-type signatures from bulk gene expression, highlighting the method’s ability to leverage gene expression programs underlying cell-type identity, even in tissues that were never observed in the training set. Using Gene Set Enrichment Analysis, we find that the metagenes learned by HYFA capture information about known biological pathways. Notably, the HYFA-imputed dataset can be used to identify regulatory genetic variations (eQTLs), with substantial gains over the original incomplete dataset. Our framework can accelerate effective and scalable integration of tissue and cell-type gene expression biorepositories.

## 1 Introduction

Sequencing technologies have enabled profiling the transcriptome at tissue and single-cell resolutions, with great potential to unveil intra- and multi-tissue molecular phenomena such as cell signalling and disease mechanisms. Due to the invasiveness of the sampling process, gene expression is usually measured independently in easy-to-acquire tissues, leading to an incomplete picture of an individual’s physiological state and necessitating effective multi-tissue integration methodologies.

A question of fundamental biological significance is to what extent the transcriptomes of difficult-to-acquire tissues and cell types can be inferred from those of accessible ones [Basu et al., 2021, Consortium, 2020]. Due to their ease of collection, accessible tissues such as whole blood could have great utility for diagnosis and monitoring of pathophysiological conditions through metabolites, signalling molecules and other biomarkers, including potential transcriptome-level associations [Yang et al., 2020]. Moreover, all human somatic cells share the same genetic information, which may regulate expression in a context-dependent and temporal manner, partially explaining tissue- and cell-type-specific gene expression variation. Computational models that exploit these patterns could therefore be used to impute the transcriptomes of uncollected cell types and tissues, with potential to elucidate the biological mechanisms regulating a diverse range of developmental and physiological processes.

Multi-tissue imputation is a central problem in transcriptomics with broad implications for fundamental biological research and translational science. The methodological problem can powerfully influence downstream applications, including performing differential expression analysis, identifying regulatory (genetic) mechanisms, determining co-expression networks, and enabling drug target discovery. In practice, in experimental follow-up or clinical application, the task includes the special case of determining a good proxy or easily-assayed system for *causal* tissues and cell types. Multi-tissue integration methods can also be applied to harmonise large collections of RNA-seq datasets from diverse institutions, consortia, and studies [Xu et al., 2021] — each potentially affected by technical artifacts — and to characterise gene expression co-regulation across tissues. Reconstruction of unmeasured gene expression across a broad collection of tissues and cell types from available reference transcriptome panels may expand our understanding of the molecular origins of complex traits and of their context specificity.

Several methods have traditionally been employed to impute uncollected gene expression. Leveraging a surrogate tissue has been widely used in studies of biomarker discovery, diagnostics, and expression Quantitative Trait Loci (eQTLs), and in the development of model systems [Hoon et al., 2000, Cai et al., 2010, Istas et al., 2017, Gamazon et al., 2018, Kim et al., 2020]. Nonetheless, gene expression is known to be tissue and cell-type specific, limiting the utility of a proxy tissue. Other related studies impute tissue-specific gene expression from genetic information [Zhou et al., 2020]. Wang et al. [2016] propose a mixed-effects model to infer uncollected data in multiple tissues from eQTLs. Sul et al. [2013] introduce a model termed Meta-Tissue that aggregates information from multiple tissues to increase statistical power of eQTL detection. However, these approaches do not model the complex, non-linear relationships between measured and unmeasured gene expression traits among tissues and cell types, and individual-level genetic information (such as at eQTLs) is subject to privacy concerns and often unavailable.

Computationally, multi-tissue transcriptome imputation is challenging because the data dimensionality scales rapidly with the number of genes and tissues, often leading to overparameterised models. TEEBoT [Basu et al., 2021] addresses this issue by employing principal component analysis (PCA) — a non-parametric dimensionality reduction method — to project the data into a low-dimensional manifold, followed by linear regression to predict target gene expression from the principal components. However, this technique does not account for non-linear effects and can handle only a single reference tissue, i.e. whole blood. Approaches such as standard multilayer perceptrons can exploit non-linear patterns, but are massively overparameterised and computationally infeasible.

To address these challenges, we present HYFA (**Hy**pergraph **Fa**ctorisation), a parameter-efficient graph representation learning approach for joint multi-tissue and cell-type gene expression imputation. HYFA represents multi-tissue gene expression in a hypergraph of individuals, metagenes, and tissues, and learns factorised representations via a specialised message passing neural network operating on the hypergraph. In contrast to existing methods, HYFA supports a variable number of reference tissues, increasing the statistical power over single-tissue approaches, and incorporates inductive biases to exploit the shared regulatory architecture of tissues. In performance comparison, HYFA attains improved performance over TEEBoT and standard imputation methods across a broad range of tissues from the Genotype-Tissue Expression (GTEx) project (v8) [Consortium, 2020]. Through transfer learning on a paired single-nucleus RNA-seq dataset (GTEx-v9) [Eraslan et al., 2021], we further demonstrate the ability of HYFA to resolve cell-type signatures from bulk gene expression. Thus, HYFA may provide a unifying transcriptomic methodology for multi-tissue imputation and cell-type deconvolution. In post-imputation analysis, application of eQTL mapping on the fully-imputed GTEx data yields a substantial increase in number of detected eQTLs. HYFA is publicly accessible at https://github.com/rvinas/HYFA.

## 2 Results

### Hypergraph factorisation: a multi-tissue transcriptomic imputation approach

We developed HYFA, a framework for inferring the transcriptomes of unmeasured tissues and cell-types from bulk expression collected in a variable number of reference tissues (Figure 1, Methods). HYFA receives as input gene expression measurements collected from a set of reference tissues, as well as demographic information, and outputs gene expression values in a tissue of interest (e.g. uncollected). The first step of the HYFA workflow is to project the input gene expression into low-dimensional *metagene* representations [Brunet et al., 2004, Raychaudhuri et al., 1999] for every collected tissue. Each metagene summarises abstract properties of groups of genes, e.g. sets of genes that tend to be expressed together [Svensson et al., 2020], that are relevant for the imputation task. In a second step, HYFA employs a custom message passing neural network [Gilmer et al., 2017] that operates on a 3-uniform hypergraph, yielding factorised individual, tissue, and metagene representations. Lastly, HYFA infers latent metagene values for the target tissue — a hyperedge-level prediction task — and maps these representations back to the original gene expression space. Through higher-order hyperedges, HYFA can also incorporate cell-type information and infer finer-grained cell-type-specific gene expression (Methods). Altogether, HYFA offers features to reuse knowledge across tissues, capture non-linear cross-tissue patterns of gene expression, learn rich representations of biological entities, and account for variable numbers of reference tissues.

**Figure 1:**
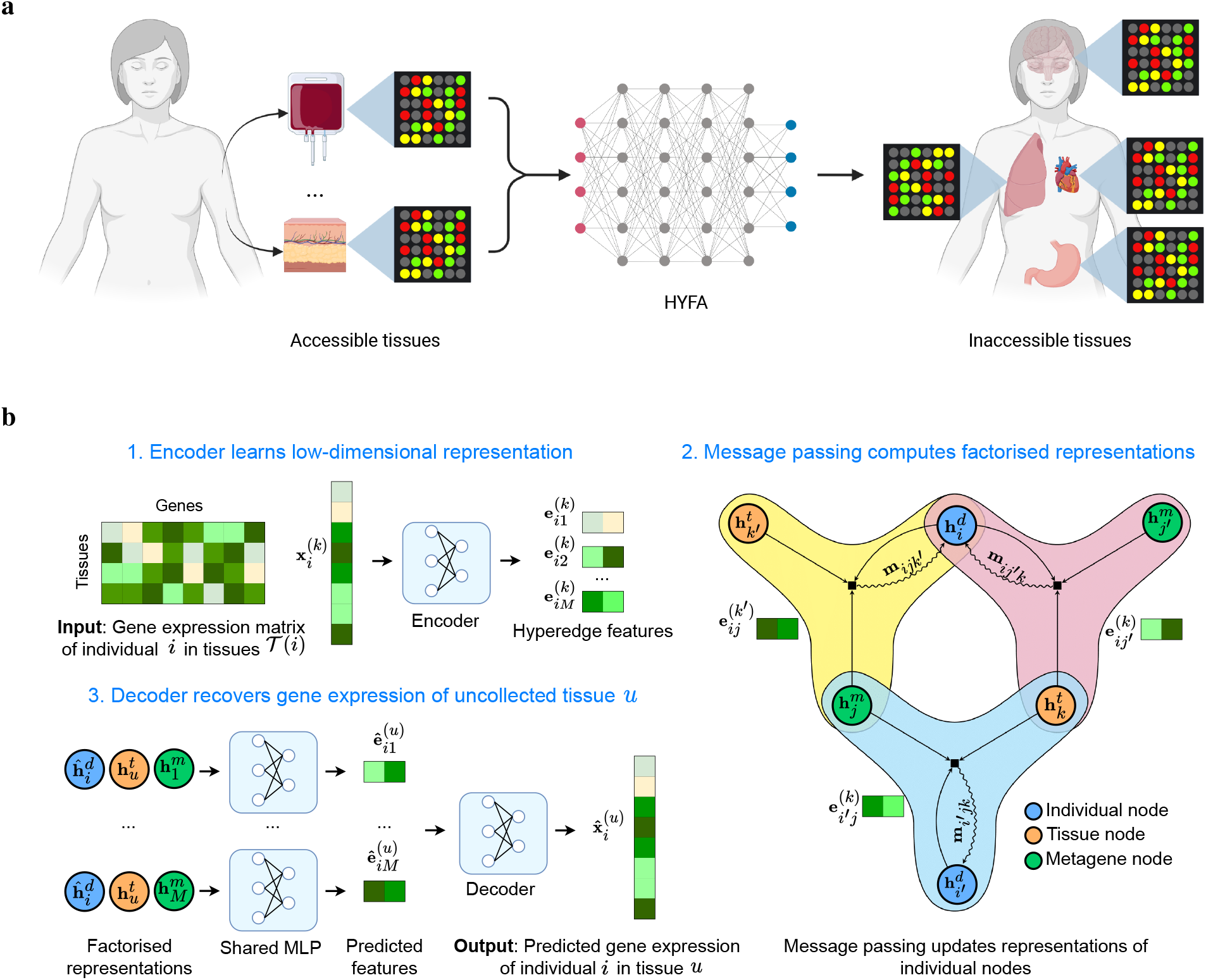
Overview of HYFA. (a) HYFA processes gene expression from a number of collected tissues (e.g. accessible tissues) and infers the transcriptomes of uncollected tissues. (b) Workflow of HYFA. The model receives as input a variable number of gene expression samples 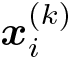 corresponding to the collected tissues *k* ∈𝒯 (*i*) of a given individual *i*. The samples 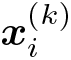 are fed through an encoder that computes low-dimensional representations 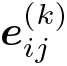 for each metagene *j* ∈ 1..*M*. A *metagene* is a latent, low-dimensional representation that captures certain gene expression patterns of the high-dimensional input sample. These representations are then used as hyperedge features in a message passing neural network that operates on a hypergraph. In the hypergraph representation, each hyperedge labelled with 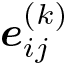 connects an individual *i* with metagene *j* and tissue *k* if tissue *k* was collected for individual *i*, i.e. *k* ∈𝒯 (*i*). Through message passing, HYFA learns factorised representations of individual, tissue, and metagene nodes. To infer the gene expression of an uncollected tissue *u* of individual *i*, the corresponding factorised representations are fed through a multilayer perceptron (MLP) that predicts low-dimensional features 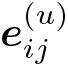 for each metagene *j* ∈ 1..*M*. HYFA finally processes these latent representations through a decoder that recovers the uncollected gene expression sample 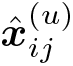.

### Characterisation of cross-tissue relationships

Characterising cross-tissue relationships at the transcriptome level can help elucidate coordinated gene regulation and expression, a fundamental phenomenon with direct implications on health homeostasis, disease mechanisms, and comorbidities [Roenneberg and Merrow, 2016, Davière and Achard, 2017, Bodine et al., 2021]. We trained HYFA on bulk gene expression from the GTEx project (GTEx-v8; Methods) [Consortium, 2020] and assessed the cross-tissue gene expression predictability scores and quality of tissue embeddings (Figure 2). The application of Uniform Manifold Approximation and Projection (UMAP) [McInnes et al., 2018] on the learnt tissue representations revealed strong clustering of biologically-related tissues (Figure 2a), including the gastrointestinal system (e.g. esophageal, stomach, colonic, and intestinal tissues), the female reproductive tissues (i.e. uterus, vagina, and ovary), and the central nervous system (i.e. the 13 brain tissues). The clustering properties were robust across UMAP runs and could be similarly appreciated using other dimensionality reduction algorithms such as t-distributed Stochastic Neighbor Embedding (t-SNE) [Van der Maaten and Hinton, 2008]. For every pair of reference and target tissues in GTEx, we then computed the Pearson correlation coefficient *ρ* between the predicted and actual gene expression, averaged the scores across individuals, and used a cutoff of *ρ* > 0.5 to depict the top pairwise associations (Figure 2b). We observed connections between most GTEx tissues and whole blood, which suggests that blood-derived gene expression is highly informative of (patho)physiological processes in other tissues [Ray et al., 2007]. Notably, brain tissues and the pituitary gland were strongly associated with several target tissues (*ρ* > 0.5), including gastrointestinal tissues (i.e. esophagus, stomach, and colon), the adrenal gland, and skeletal muscle, which may account for known disease comorbidities.

**Figure 2:**
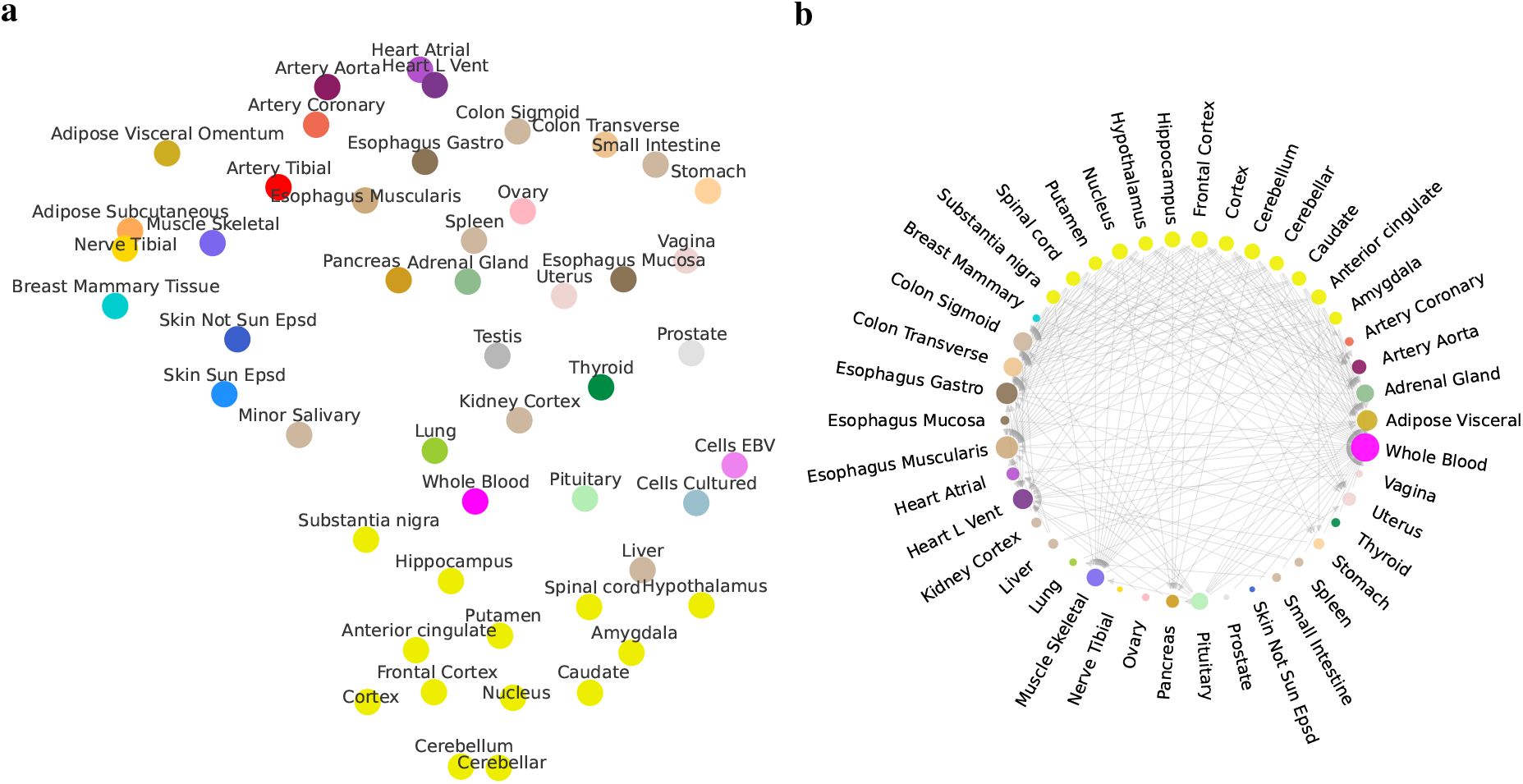
Analysis of cross-tissue relationships. Colors are assigned to conform to the GTEx Consortium conventions. (a) UMAP representation of the tissue embeddings learnt by HYFA. Note that human body systems cluster in the embedding space (e.g. digestive system: stomach, small intestine, colon, esophagus; and central nervous system). (b) Network of tissues depicting the predictability of target tissues with HYFA. The dimension of each node is proportional to its degree. Edges from reference to target tissues indicates an average Pearson correlation coefficient *ρ* > 0.5. Interestingly, central nervous system tissues strongly correlate with several non-brain tissues such as gastrointestinal tissues and skeletal muscles.

### Predictability of tissue-specific gene expression from whole blood transcriptome

Knowledge about tissue-specific patterns of gene expression can increase our understanding of disease biology, facilitate the development of diagnostic tools, and improve patient subtyping [Lage et al., 2008, Basu et al., 2021], but most tissues are inaccessible or difficult to acquire. To address this challenge, we studied to what extent HYFA can recover tissue-specific gene expression from whole-blood transcriptomic measurements (Figures 3a and 3c). For each individual in the test set with measured whole-blood gene expression, we employed the hypergraph neural network to predict tissue-specific gene expression in the individual’s remaining collected tissues. We evaluated imputation performance using the Pearson correlation coefficient *ρ* between the high-dimensional inferred gene expression and the ground-truth target samples. We observed strong prediction scores for esophageal tissues (muscularis: *ρ* = 0.488, gastro: *ρ* = 0.461, mucosa: *ρ* = 0.356), heart tissues (left ventricle: *ρ* = 0.479, atrial: *ρ* = 0.455) and lung (*ρ* = 0.471), while Epstein Barr virus-transformed lymphocytes (*ρ* = 0.057), an accessible and renewable resource for functional genomics, was a notable outlier. We compared our method with TEEBoT [Basu et al., 2021] (without expression single-nucleotide polymorphism information), which first projects the high-dimensional blood expression data into a low-dimensional space through principal component analysis (30 components; 75-80% explained variance) and then performs linear regression to predict the gene expression of the target tissue. Overall, TEEBoT and HYFA attained comparable scores when a single tissue (i.e. whole blood) was used as reference and both methods outperformed standard imputation approaches (mean imputation, blood surrogate, and k nearest neighbours; Figure 3c).

**Figure 3:**
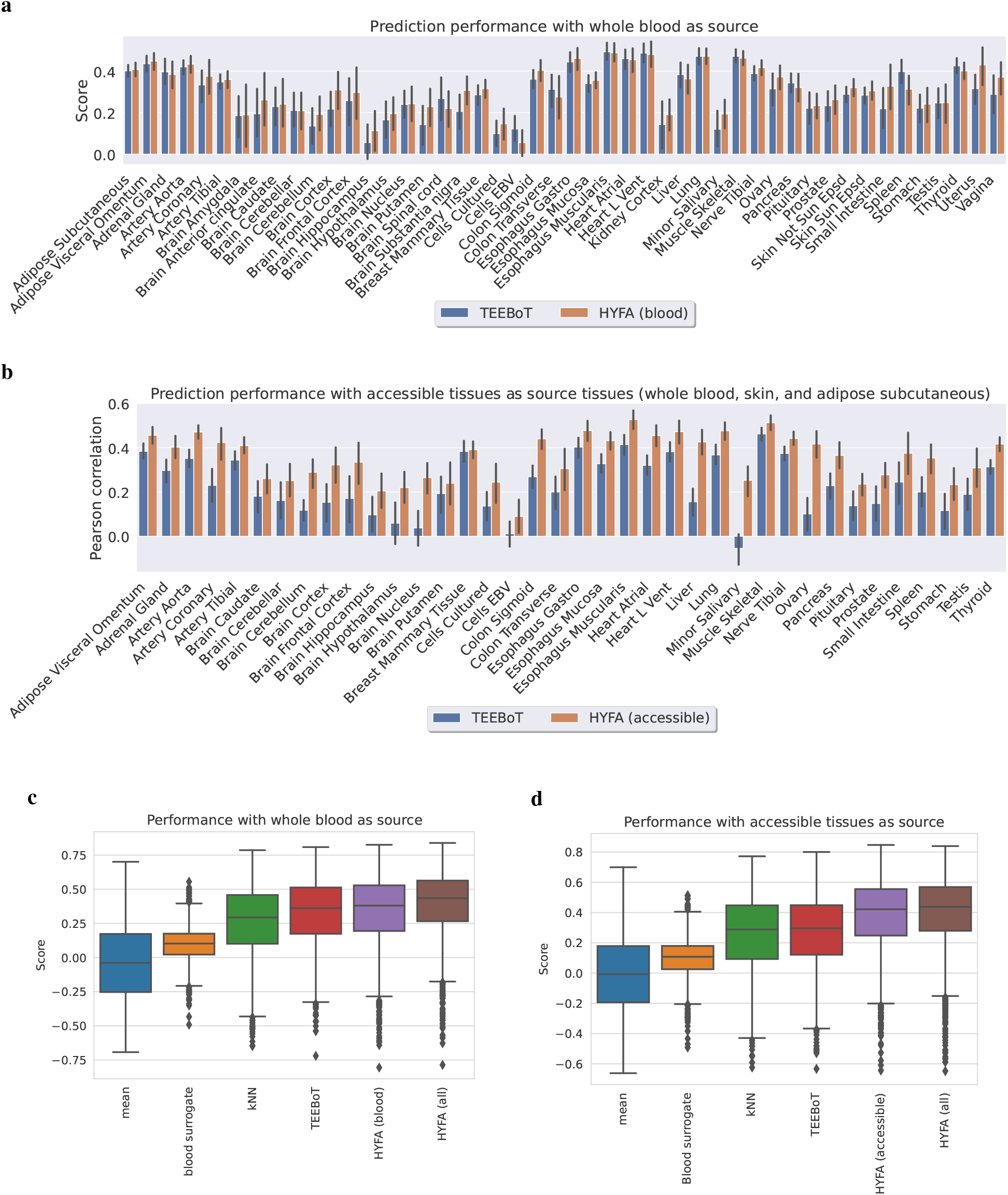
Performance comparison across gene expression imputation methods. Performance comparison across gene expression imputation methods. Error bars indicate 95% confidence intervals. (a, b) Per-tissue comparison between HYFA and TEEBoT when using (a) whole-blood and (b) all accessible tissues (whole blood, skin sun exposed, skin not sun exposed, and adipose subcutaneous) as reference. We discarded target tissues represented by less than 25 test individuals. HYFA achieved superior Pearson correlation in (a) 25 out of 48 target tissues when a single tissue was used as reference and (b) all target tissues when multiple reference tissues were considered. (c, d) Prediction performance from (c) whole-blood gene expression and (d) accessible tissues as reference. Mean imputation replaces missing values with the feature averages. Blood surrogate utilises gene expression in whole blood as a proxy for the target tissue. k-Nearest Neighbours (kNN) imputes missing features with the average of measured values across the k nearest observations (k=20). TEEBoT projects reference gene expression into a low-dimensional space with principal component analysis (PCA; 30 components), followed by linear regression to predict target values. HYFA (all) employs information from all collected tissues.

### Multiple reference tissues improve performance

We hypothesised that using multiple tissues as reference would improve downstream imputation performance. To evaluate this, we selected individuals with measured gene expression both at the target tissue and 4 reference (accessible) tissues (whole blood, skin sun exposed, skin not sun exposed, and adipose subcutaneous) and employed HYFA to impute target expression values (Figure 3 and Extended Data Figure 1). We discarded underrepresented target tissues with less than 25 test individuals. Relative to using whole blood in isolation, using all accessible tissues as reference resulted in improved performance for 14 out of 18 target tissues (Extended Data Figure 1), highlighting the benefit of leveraging multiple reference tissues. This particularly boosted imputation scores for esophageal tissues (muscularis: Δ*ρ* = 0.068, gastro: Δ*ρ* = 0.061, mucosa: Δ*ρ* = 0.048), colonic tissues (transverse: Δ*ρ* = 0.065, sigmoid: Δ*ρ* = 0.056), and artery tibial (Δ*ρ* = 0.079) (Extended Data Figure 1). In contrast, performance for the pituitary gland (Δ*ρ* = −0.011), lung (Δ*ρ* = −0.003), and stomach (Δ*ρ* = −0.002) remained stable or dropped slightly. Moreover, the performance gap between HYFA and TEEBoT (trained on the set of complete multi-tissue samples) further widened relative to the single-tissue scenario (Figure 3) — HYFA obtained better performance in all target tissues. We attribute the improved scores to HYFA’s ability to process a variable number of reference tissues, reuse knowledge across tissues, and capture non-linear patterns, resulting in increased statistical strength and better generalisation to unseen combinations of reference-target tissues.

### Inference of cell-type signatures

We next investigated the ability of HYFA to predict cell-type-specific signatures in a given tissue of interest. We first matched donors with collected bulk gene expression in GTEx-v8 Consortium [2020] and individuals with measured single-nucleus RNA-seq profiles from GTEx-v9 [Eraslan et al., 2021], resulting in 16 individuals. We selected the top 3000 highly variable genes using the Scanpy function `scanpy.pp.highly_variable_genes` with flavour setting `seurat_v3` [Wolf et al., 2018, Stuart et al., 2019] (Methods). We then aggregated the snRNA-seq profiles for each cell-type, tissue, and individual in GTEx-v9, yielding 226 unique individual-specific signatures. We discarded under-represented cell-types with less than 10 signatures and split the signatures into train and test (Extended Data Figure 2). Next, we trained HYFA to infer the cell-type, tissue, and individual-specific signatures from the multi-tissue bulk expression profiles. We evaluated performance using the observed (Figure 2) and inferred library sizes (Extended Data Figure 3). To attenuate the small data size problem, we applied transfer learning on the model trained for the multi-tissue imputation task (Methods). We observed strong prediction scores for vascular endothelial cells (heart: *ρ* = 0.79; breast: *ρ* = 0.87, esophagus muscularis: *ρ* = 0.79) and fibroblasts (heart: *ρ* = 0.80; breast: *ρ* = 0.89, esophagus muscularis: *ρ* = 0.80). Strikingly, HYFA recovered the cell-type profiles of tissues that were never observed in the train set with high correlation (Figures 2 and 4), e.g. skeletal muscle (vascular endothelial cells: *ρ* = 0.80, fibroblasts: *ρ* = 0.76, pericytes/SMC: *ρ* = 0.70), demonstrating the benefits of the transfer learning approach and the factorised tissue representations. Our results highlight the potential of HYFA to impute unknown cell-type signatures even for tissues that were not considered in the original snRNA-seq dataset [Eraslan et al., 2021].

**Figure 4:**
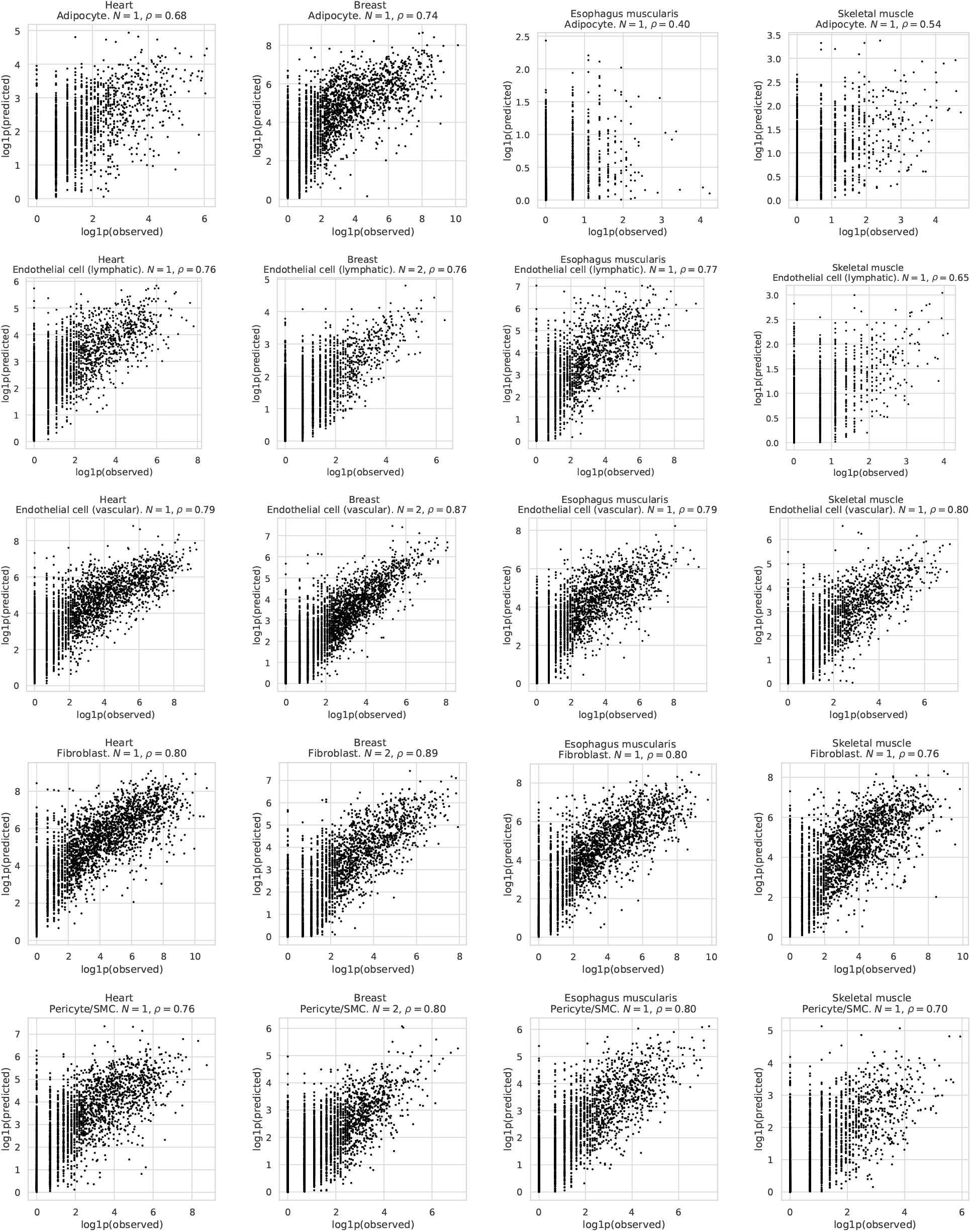
Prediction of cell-type signatures. HYFA imputes individual- and tissue-specific cell-type signatures from bulk multi-tissue gene expression. The scatter plots depict the Pearson correlation *ρ* between the logarithmised ground truth and predicted signatures for *N* unseen individuals. To infer the signatures, we used the observed library size 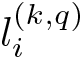 and number of cells 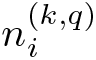.

### Multi-tissue imputation improves eQTL detection

Gene expression acts as an intermediate molecular trait between DNA and phenotype and, therefore, genetic mapping of genome-wide gene expression can shed light on the genetic architecture and molecular basis of complex traits. The GTEx project has enabled the identification of numerous genetic associations with gene expression across a broad collection of tissues [Consortium, 2020], also known as expression Quantitative Trait Loci (eQTLs) [Nica and Dermitzakis, 2013]. However, eQTL datasets are characterised by small sample sizes, especially for difficult-to-acquire tissues and cell types, reducing the statistical power to detect eQTLs [Rockman and Kruglyak, 2006]. To address this problem, we employed HYFA to impute the transcript levels of every uncollected tissue for each individual in GTEx, yielding a complete gene expression dataset of 834 individuals and 49 tissues. We then performed eQTL mapping (Methods) on the original and imputed datasets and observed a substantial gain in the number of unique genes with detected eQTLs (Figure 5). Notably, this metric increased for tissues with low sample size (Spearman correlation coefficient *ρ* = −0.83) — which are most likely to benefit from borrowing information across tissues with shared regulatory architecture. Kidney cortex displayed the largest gain (from 215 to 12,557), while there was no increase observed for whole blood. Our results generate a large catalog of new tissue-specific eQTLs (Data availability), with potential to enhance our understanding of how regulatory variation mediates variation in complex traits, including disease susceptibility.

**Figure 5:**
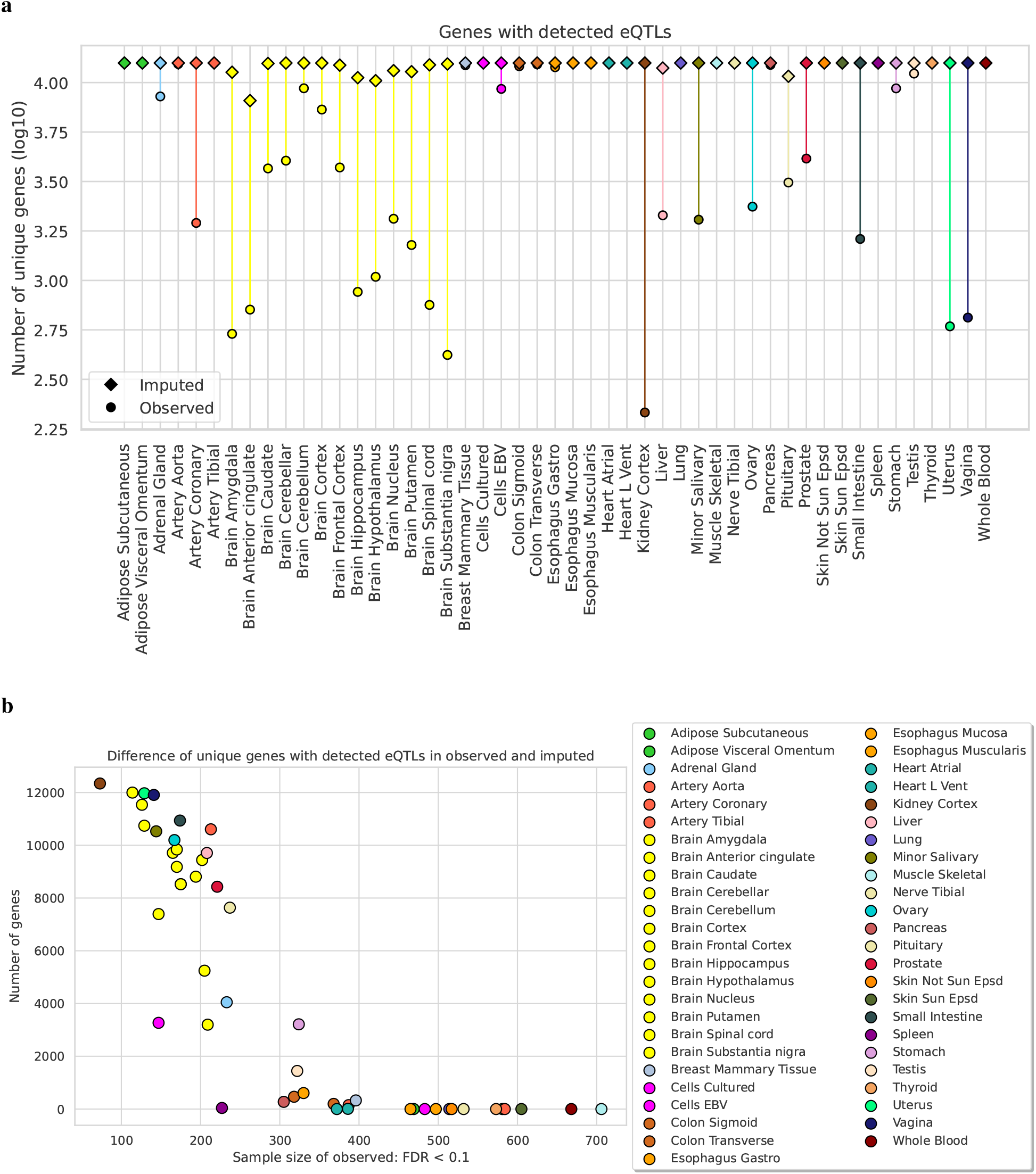
HYFA’s imputed data improves expression Quantitative Trait Loci (eQTL) discovery. (a) Number of unique genes with detected eQTLs (FDR < 0.1) on observed (circle) and full (observed plus imputed; rhombus) GTEx data. Note logarithmic scale of y-axis. (b) Increase in number of unique genes with mapped eQTLs as a function of observed sample size. HYFA’s imputed data substantially increases the number of identified associations, especially for tissues with low sample sizes.

### Brain-gut axis

The brain-gut axis is a bidirectional communication system of signalling pathways linking the central and enteric nervous systems. We investigated the extent to which the transcriptomes of tissues from the gastrointestinal system are predictive of gene expression in brain tissues. We selected all the unseen individuals with simultaneous measurements in gastrointestinal tissues (i.e. oesophago gastric junction) and brain tissues (i.e. frontal cortex, hippocampus, and anterior cingulate) and employed HYFA to predict the expression values of brain tissues (Figure 6). We observed a small number of individuals with measurements in both brain and non-brain tissues (Supplementary Materials D). After ranking the genes according to their predicability scores and selecting the top 1000 genes for each brain tissue (Venn diagram; Figure 6), we found considerable overlap between the 3 brain tissues (153 common genes in the intersection). We then used Enrichr [Kuleshov et al., 2016] with the gene sets *GO_Biological_Process_2021* and *GO_Molecular_Function_2021* to identify the enriched Gene Ontology (GO) terms for the shared genes at the intersection. Overall, the best predicted genes were enriched in multiple signalling-related terms (e.g. cytokine receptor activity and interleukin-1 receptor activity). This aligns with studies that highlight that gut microbes communicate with the central nervous system through endocrine and immune signalling mechanisms [Martin et al., 2018]. Genes in the intersection were also notably enriched in the ciliary neurotrophic factor receptor activity (molecular function), which plays an important role in the survival of neurons [Davis et al., 1991], the development of the enteric nervous system [Liu, 2018], and the control of body weight [Xu and Xie, 2016]. Moreover, our results suggest an association with the Receptor for Advanced Glycation Endproducts (RAGE), which has been linked to inflammation-related pathological states such as vascular disease, diabetes, and neurodegeneration [Sparvero et al., 2009].

**Figure 6:**
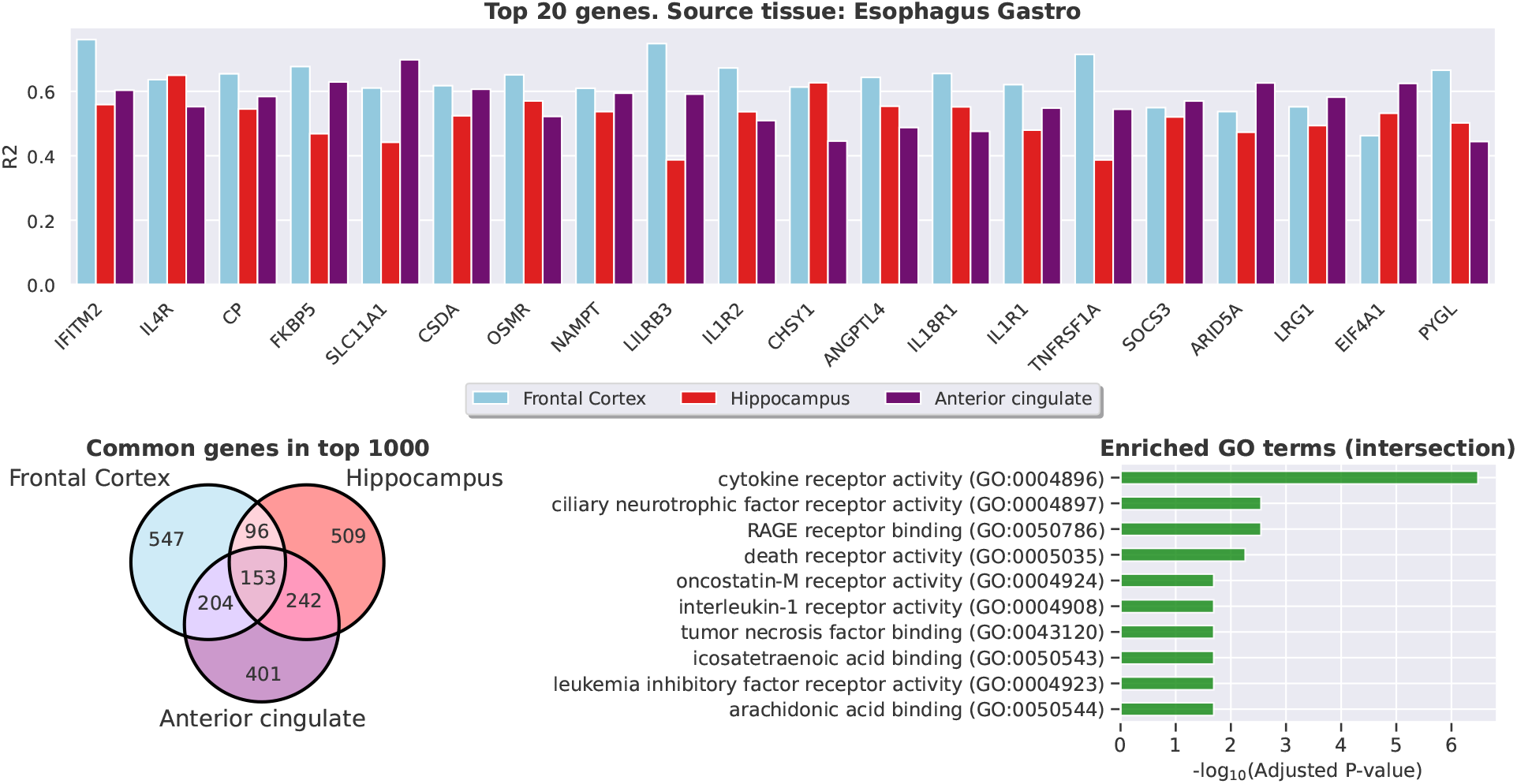
Top predicted genes in multiple brain regions with the oesophago gastric junction as the reference tissue. (a) ℝ^2^ coefficient of top 20 predicted genes, ranked by average score. (b) Common genes in the top 1000 predicted genes for each brain tissue. (c) Enriched GO terms of the top shared genes at the interesection. The best predicted genes were enriched in signalling pathways (FDR < 0.05), consistent with studies reporting that gut microbes communicate to the central nervous system through endocrine and immune mechanisms. Moreover, these results highlight the potential association between the elements of the oesophago gastric junction and the ciliary neurotrophic factor, which has been linked to the survival of neurons [Davis et al., 1991] and the control of body weight [Xu and Xie, 2016].

### HYFA-learned metagenes capture known biological pathways

A key feature of HYFA is that it reuses knowledge across tissues and metagenes, allowing to exploit shared regulatory patterns. We explored whether HYFA’s inductive biases encourage learning biologically relevant metagenes. To determine the extent to which metagene-factors relate to known biological pathways, we applied Gene Set Enrichment Analysis (GSEA) [Subramanian et al., 2005] to the gene loadings of HYFA’s encoder (Methods). Similar to [Zhao et al., 2021], for a given query gene set, we calculated the maximum running sum of enrichment scores by descending the sorted list of gene loadings for every metagene and factor. We then computed pathway enrichment p-values through a permutation test and employed the Benjamini-Hochberg method to correct for multiple testing. In total, we identified 18683 statistically significant enrichments (FDR < 0.05) of KEGG biological processes across all HYFA metagenes and factors (Figure 7a). Among these, 2109 corresponded to signalling pathways and 1300 to pathways of neurodegeneration (Figure 7b). We observed considerable overlap between several metagenes in terms of enriched biologically related pathways, e.g. factor 95 of metagene 11 had the lowest FDR for both Alzheimer’s disease (FDR < 0.001) and Amyotrophic Lateral Sclerosis (FDR < 0.001) pathways (Figure 7d and 7f). We then applied UMAP to project the latent values of metagene 11 and observed relatively strong clustering of individuals with Amyotrophic Lateral Sclerosis (spinal cord; Figure 7c) and Alzheimer’s disease or Dementia (brain cortex, Figure 7e). Altogether, our analysis suggests that HYFA-learned metagenes capture information about known pathways and are therefore amenable to biological interpretation.

**Figure 7:**
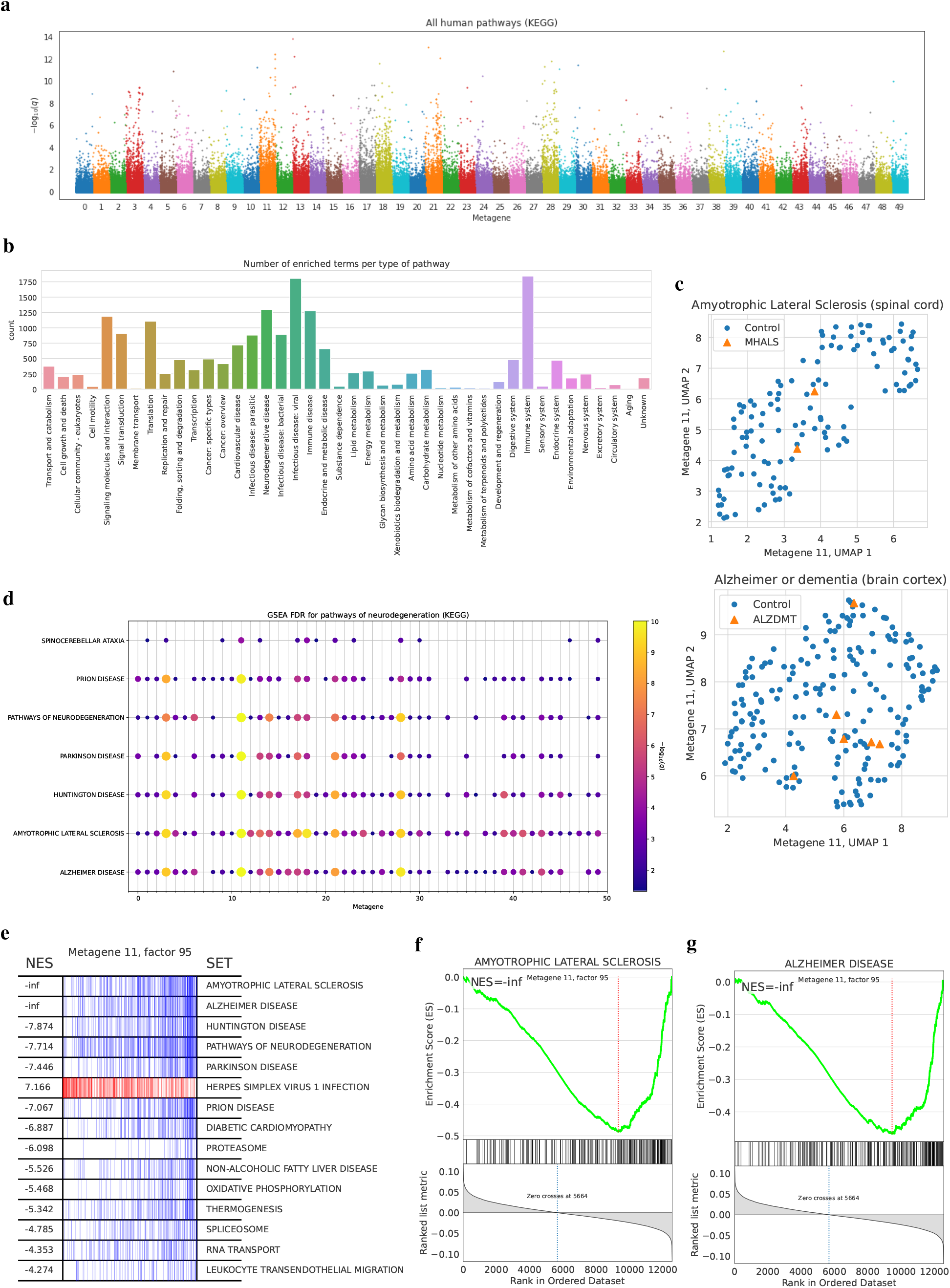
Pathway enrichment analysis of metagene factors. Pathway enrichment analysis of metagene factors. (a) Manhattan plot of the GSEA results on the metagenes (n=50) and factors (n=98) learned by HYFA. The x-axis represents metagenes (colored bins) and every offset within the bin corresponds to a different factor. The y-axis is the –log q-value (FDR) from the GSEA permutation test, corrected for multiple testing via the Benjamini-Hochberg procedure. We identified 18683 statistically significant enrichments (FDR < 0.05) of KEGG biological processes across all metagenes and factors. (b) Total number of enriched terms for each type of pathway. (d) FDR for pathways of neurodegeneration. For every pathway and metagene, we selected the factor with lowest FDR and depicted statistically significant values (FDR < 0.05). Point sizes are proportional to − log FDR values. Metagene 11 (factor 95) had the lowest FDR for both Amyotropic Lateral Sclerosis (ALS) and Alzheimer’s Disease. (c) UMAP of latent values of metagene 11 for all spinal cord (c, ALS: orange) and brain cortex (e, Alzheimer’s Disease or Dementia: orange) GTEx samples. (e) Leading edge subsets of top-15 enriched gene sets for factor 95 of metagene 11. (f, g) Enrichment plots for Amyotropic Lateral Sclerosis (f) and Alzheimer’s disease gene sets (g).

## 3 Discussion

Effective multi-tissue omics integration promises a system-wide view of human physiology, with potential to shed light on intra- and multi-tissue molecular phenomena. Such an approach challenges single-tissue and conventional integration approaches — often unable to model a variable number of tissues with sufficient statistical strength — necessitating the development of scalable, non-linear, and flexible methods. Here we developed HYFA (**Hy**pergraph **Fa**ctorisation), a parameter-efficient approach for joint multi-tissue and cell-type gene expression imputation that imposes strong inductive biases to learn entity-independent relational semantics and demonstrates excellent imputation capabilities. The hypergraph factorisation framework is flexible (it supports k-uniform hypergraphs) and may find application beyond computational genomics.

We performed extensive benchmarks on GTEx (v8) data [Consortium, 2020], the most comprehensive human transcriptome resource available, and evaluated imputation performance over a broad collection of tissues and cell types. In addition to standard transcriptome imputation approaches, we compared our method with TEEBoT [Basu et al., 2021], a linear method that predicts target gene expression from the principal components of the reference expression. In the single-tissue reference scenario, HYFA and TEEBoT attained comparable imputation scores, outperforming standard methods. In the multi-tissue reference scenario, HYFA consistently outperformed TEEBoT and standard approaches for all target tissues, demonstrating the effectiveness of HYFA’s capabilities to borrow non-linear information across a variable number of tissues and exploit shared molecular patterns.

In addition to imputing tissue-level transcriptomics, we investigated the ability of HYFA to predict cell-type-level gene expression from multi-tissue bulk expression measurements. Through transfer learning, we trained HYFA to infer cell-type signatures from a cohort of single-nucleus RNA-seq [Eraslan et al., 2021] with matching GTEx-v8 donors. The inferred cell-type signatures exhibited a strong correlation with the ground truth despite the low sample size, indicating that HYFA’s latent representations are rich and amenable to knowledge transfer. Strikingly, HYFA also recovered cell-type profiles from tissues that were never observed at transfer time, pointing to HYFA’s ability to leverage gene expression programs underlying cell-type identity [Kotliar et al., 2019] even in tissues that were not considered in the original study of Eraslan et al. [2021]. For future work, we will investigate extensions of HYFA to resolve the cell-type composition of observed and imputed tissue-specific gene expression.

In post-imputation analysis, we studied whether the imputed data improves eQTL discovery. We employed HYFA to impute the gene expression levels of every uncollected tissue in GTEx-v8, yielding a complete dataset, and performed eQTL mapping. Compared to the original dataset, we observed a substantial gain in number of genes with detected eQTLs, with kidney cortex showing the largest gain. The increase was highest for tissues with low sample sizes, which are the ones expected to benefit the most from knowledge sharing across tissues. Our results uncover a large number of previously undetected tissue-specific eQTLs and highlight the ability of HYFA to exploit shared regulatory information across tissues.

In addition to imputing uncollected gene expression measurements, HYFA can provide insights on coordinated gene regulation and expression mechanisms across tissues. We analysed to what extent tissues from the gastrointestinal system are informative of gene expression in brain tissues — an important question that may shed light on the biology of the brain-gut axis — and identified enriched biological processes and molecular functions. Through Gene Set Enrichment Analysis [Subramanian et al., 2005], we further introspected the HYFA-learned metagenes and found a substantial amount of enriched pathways, opening the door to further biological interpretations. We believe that explainability methods could be integrated in the HYFA pipeline to generate interesting biological insights. Future work might also seek to impose stronger inductive bias to ensure that metagenes are identifiable and robust to batch effects.

We believe that HYFA, as a versatile graph representation learning framework, provides a novel methodology for effective integration of large-scale multi-tissue biorepositories. HYFA’s implementation is available online at https://github.com/rvinas/HYFA.

## 4 Methods

### 4.1 Problem formulation

Suppose we have a transcriptomics dataset of *N* individuals/donors, *T* tissues, and *G* genes. For each individual *i* ∈ {1, &, *N*}, let ***X***_*i*_ ∈ ℝ^*T* ×*G*^ be the gene expression values in *T* tissues and define the donor’s demographic information by ***u***_*i*_ ∈ ℝ^*C*^, where *C* is the number of covariates. Denote by 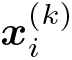 the *k*-th entry of ***X***_*i*_, corresponding to the expression values of donor *i* measured in tissue *k*. For a given donor *i*, let 𝒯(*i*) represent the collection of tissues with measured expression values. These sets might vary across individuals. Let 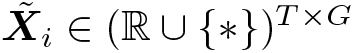 be the measured gene expression values, where * denotes unobserved, so that 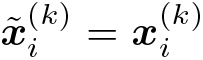 if *k* ∈𝒯(*i*) and 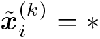 otherwise. Our goal is to infer the uncollected values in 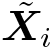 by modelling the distribution 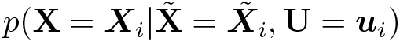.

### 4.2 Multi-tissue model

An important challenge of modelling multi-tissue gene expression is that a different set of tissues might be collected for each individual. Moreover, the data dimensionality scales rapidly with the total number of tissues and genes. To address these problems, we represent the data in a hypergraph and develop a parameter-efficient neural network that operates on this hypergraph. Throughout, we make use of the concept of *metagenes* [Brunet et al., 2004, Raychaudhuri et al., 1999]. Each *metagene* characterises certain gene expression patterns and is defined as a positive linear combination of multiple genes [Brunet et al., 2004, Raychaudhuri et al., 1999].

#### 4.2.1 Hypergraph representation

We represent the data in a hypergraph consisting of three types of nodes: donor, tissue, and *metagene* nodes. Mathematically, we define a hypergraph 𝒢 = {𝒱_*d*_ ∪𝒱_*m*_ ∪𝒱_*t*_, ℰ}, where 𝒱_*d*_ is a set of donor nodes, 𝒱_*m*_ is a set of *metagene* nodes, 𝒱_*t*_ is a set of tissue nodes, and ℰ is a set of multi-attributed hyperedges. Each hyperedge connects an individual *i* with a *metagene j* and a tissue *k* if *k* ∈𝒯 (*i*), where 𝒯 (*i*) are the collected tissues of individual *i*. The set of all hyperedges is defined as, 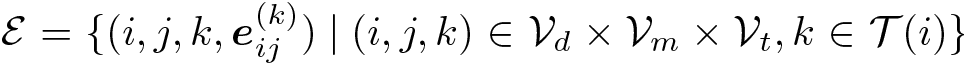, where 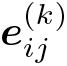 are hyperedge attributes that describe characteristics of the interacting nodes, i.e. features of *metagene j* in tissue *k* for individual *i*. The hypergraph representation allows representing data in a flexible way, generalising the bipartite graph representation from You et al. [2020]. On the one hand, using a single *metagene* results in a bipartite graph where each edge connects an individual *i* with a tissue *k*. In this case, the edge attributes 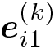 are derived from the gene expression 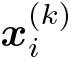 of individual *i* in tissue *k*. On the other hand, using multiple *metagenes* leads to a hypergraph where each individual *i* is connected to tissue *k* through multiple hyperedges. For example, it is possible to construct a hypergraph where genes and *metagenes* are related by a one-to-one correspondence, with hyperedge attributes 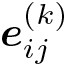 derived directly from expression 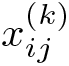. The number of *metagenes* thus controls a spectrum of hypergraph representations and, as we shall see, can help alleviate the inherent over-squashing problem of graph neural networks.

#### 4.2.2 Message passing neural network

Given the hypergraph representation of the multi-tissue transcriptomics dataset, we now present a parameter-efficient graph neural network to learn donor, metagene, and tissue embeddings, and infer the expression values of the unmeasured tissues. We start by computing hyperedge attributes from the multi-tissue expression data. Then, we initialise the embeddings of all nodes in the hypergraph, construct the message passing neural network, and define an inference model that builds on the latent node representations obtained via message passing.

##### Computing hyperedge attributes

We first reduce the dimensionality of the measured transcriptomics values. For every individual *i* and measured tissue *k*, we project the corresponding gene expression values 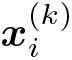 into low-dimensional *metagene* representations 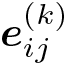:

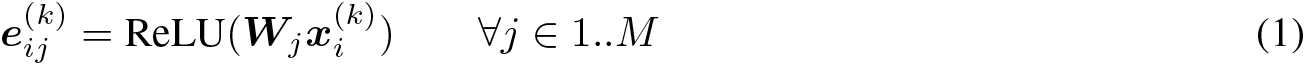

where *M*, the number of metagenes, is a user-definable hyperparameter and ***W***_*j*_ ∀*j* ∈ 1..*M* are learnable parameters. In addition to characterising groups of functionally similar genes, employing metagenes reduces the number of messages being aggregated for each node, addressing the over-squashing problem of graph neural networks (Supplementary Materials H).

##### Initial node embeddings

We initialise the node features of the individual 𝒱_*p*_, metagene 𝒱_*m*_, and tissue 𝒱_*t*_ partitions with learnable parameters and available information. For metagene and tissue nodes, we use learnable embeddings as initial node values. The idea is that these weights, which will be approximated through gradient descent, should summarise relevant properties of each metagene and tissue. We initialise the node features of each individual with the available demographic information ***u***_*i*_ of each individual *i* (e.g. age and sex). Importantly, this formulation allows transfer learning between sets of distinct donors.

##### Message passing layer

We develop a custom GNN layer to compute latent donor embeddings by passing messages along the hypergraph. At each layer of the GNN, we perform message passing to iteratively refine the individual node embeddings. We do not update the tissue and metagene embeddings during message passing, in a similar vein as knowledge graph embeddings [Bordes et al., 2013], because their node embeddings already consist of learnable weights that are updated through gradient descent. Sending messages to these nodes would also introduces a dependency between individual nodes and tissue and metagene features (and, by transitivity, dependencies between individuals). However, if we foresee that unseen entities will be present at test time (e.g. new tissue types), our approach can be extended by initialising their node features with constant values and introducing node-type-specific message passing equations.

Mathematically, let 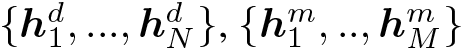, and 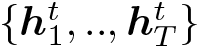 be the donor, metagene, and tissue node embeddings, respectively. At each layer of the GNN, we compute refined individual embeddings 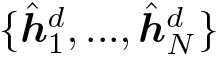 as follows:

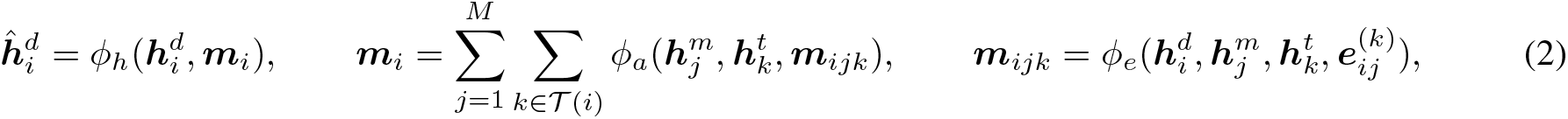

where the functions *ϕ*_*e*_ and *ϕ*_*h*_ are edge and node operations that we model as multilayer perceptrons (MLP), and *ϕ*_*a*_ is a function that determines the aggregation behavior. In its simplest form, choosing 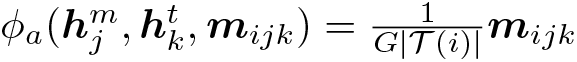 results in average aggregation. Optionally, we can stack several message passing layers to increase the expressivity of the model.

The architecture is flexible to allow the following extensions:

- Incorporation of information about the individual embeddings 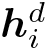 into the aggregation mechanism *ϕ*_*a*_.
- Incorporation of target tissue embeddings 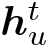, for a given target tissue *u*, into the aggregation mechanism *ϕ*_*a*_.
- Update hyperedge attributes 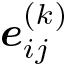 at every layer.

##### Aggregation mechanism

In practice, the proposed hypergraph neural network suffers from a bottleneck. In the aggregation step, the number of messages being aggregated is *M*|𝒯 (*i*)| for each individual *i*. In the worst case, when all genes are used as metagenes (i.e. *M* = *G*; it is estimated that humans have around *G* ≈ 25, 000 protein-coding genes), this leads to serious over-squashing — large amounts of information are compressed into fixed-length vectors [Alon and Yahav, 2020]. Fortunately, choosing a small number of metagenes reduces the dimensionality of the original transcriptomics values which in turn alleviates the over-squashing and scalability problems (Supplementary Materials H). To further attenuate over-squashing, we propose an attention-based aggregation mechanism *ϕ*_*a*_ that weighs metagenes according to their relevance in each tissue:

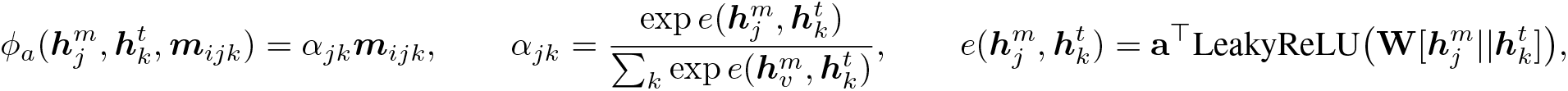

where || is the concatenation operation and **a** and **W** are learnable parameters. The proposed attention mechanism, which closely follows the neighbour aggregation method of graph attention networks [Brody et al., 2022, Veličković et al., 2018], computes dynamic weighting coefficients that prioritise messages coming from important metagenes. Optionally, we can leverage multiple heads [Vaswani et al., 2017] to learn multiple modes of interaction and increase the expressivity of the model.

#### Hypergraph model

The hypergraph model, which we define as *f*, computes latent individual embeddings 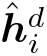 from incomplete multi-tissue expression profiles as 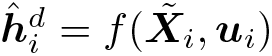

### 4.3 Downstream imputation tasks

The resulting donor representations 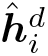 summarise information about a variable number of tissue types collected for donor *i*, in addition to demographic information. We leverage these embeddings for two downstream tasks: inference of gene expression in uncollected tissues and prediction of cell-type signatures.

#### 4.3.1 Inference of gene expression in uncollected tissues

Predicting the transcriptomic measurements 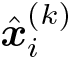 of a tissue *k* (e.g. uncollected) is achieved by first recovering the latent metagene values 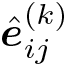 for all metagenes *j* ∈ 1..*M*, a hyperedge-level prediction task, and then decoding the gene expression values from the predicted metagene representations 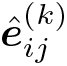 with an appropriate probabilistic model.

##### Prediction of hyperedge attributes

To predict the latent metagene attributes 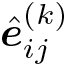 for all *j* ∈ 1..*M*, we employ a *factorised* metagene 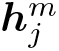 and tissue representations 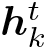 as well as the latent multilayer perceptron that operates on the variables 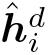 of individual *i*:

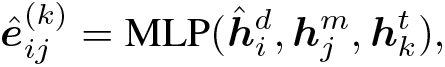

where the MLP is shared for all combinations of metagenes, individuals, and tissues.

##### Negative binomial imputation model

For raw count data, we use a negative binomial likelihood. To decode the gene expression values for a tissue *k* of individual *i*, we define the following probabilistic model 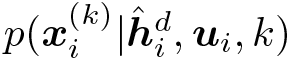:

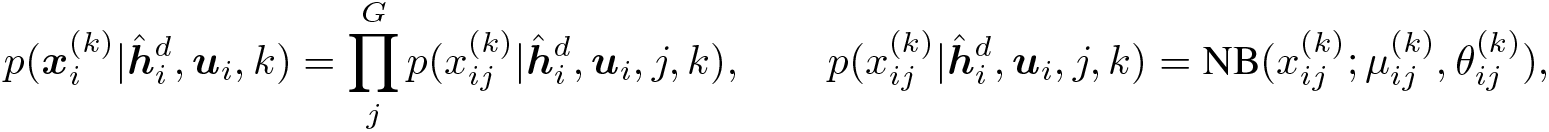

where NB is a negative binomial distribution. The mean 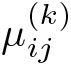 and dispersion 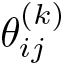 parameters of this distribution are computed as follows:

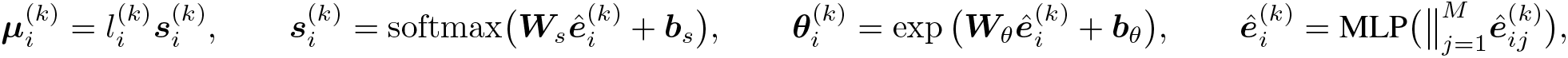

where 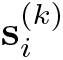 are mean gene-wise proportions; ***W***_*s*_, ***W***_*θ*_, ***b***_*s*_, and ***b***_*θ*_ are learnable parameters; and 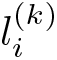 is the library size, which is modelled with a log-normal distribution

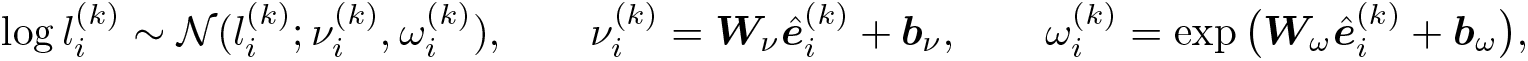

where ***W***_*ν*_, ***W***_*ω*_, ***b***_*ν*_, and ***b***_*ω*_ are learnable parameters. Optionally, we can use the observed library size.

##### Gaussian imputation model

For normalised gene expression data (i.e. inverse normal transformed data), we use the following Gaussian likelihood:

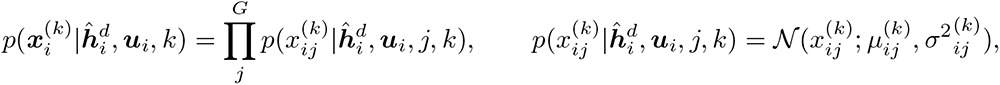

where the mean 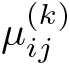 and standard deviation 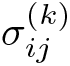 are computed as follows:

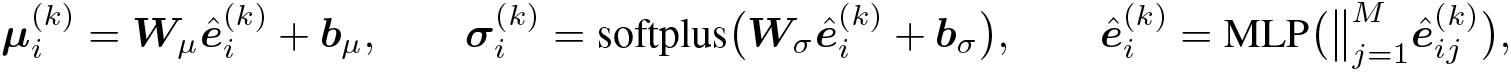

where ***W***_*μ*_, ***W***_*σ*_, ***b***_*μ*_, and ***b***_*σ*_ are learnable parameters and softplus(*x*) = log (1 + exp(*x*)).

##### Optimisation

We optimise the model to maximise the imputation performance on a dynamic subset of observed tissues, that is, tissues that are masked out at train time, similar to Viñas et al. [2021]. For each individual *i*, we randomly select a subset 𝒞 ⊂𝒯 (*i*) of *pseudo-observed* tissues and treat the remaining tissues 𝒰 = 𝒯(*i*) −𝒞 as unobserved (*pseudo-missing*). We then compute the individual embeddings 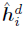 using the gene expression of *pseudo-observed* tissues 𝒞 and minimise the loss:3

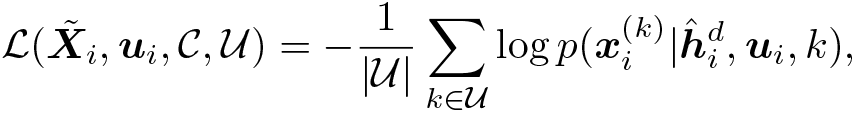

which corresponds to the average negative log-likelihood across *pseudo-missing* tissues. Importantly, the *pseudo-mask* mechanism generates different sets of *pseudo-missing* tissues for each individual, effectively enlarging the number of training examples and regularising our model. We summarise the training algorithm in Algorithm 1 (Supplementary Materials F).

##### Inference of gene expression from uncollected tissues

At test time, we infer the gene expression values 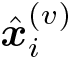 of an uncollected tissue *v* from a given donor *i* via the mean, i.e. 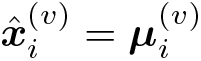. Alternatively, we can draw random samples from the conditional predictive distribution 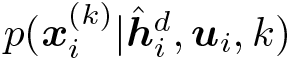.

#### 4.3.2 Prediction of cell-type signatures

We next consider the problem of imputing cell-type signatures in a tissue of interest. We define a cell-type signature as the sum of gene expression profiles across cells of a given cell-type in a certain tissue. Formally, let 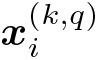 be the gene expression signature of cell-type *q* in a tissue of interest *k* of individual *i*. Our goal is to infer 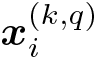 from the multi-tissue gene expression measurements 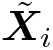. To achieve this, we first compute the hyperedge features of a hypergraph consisting of 4-node hyperedges and then infer the corresponding signatures with a zero-inflated model.

##### Prediction of hyperedge attributes

We consider a hypergraph where each hyperedge groups an individual, a tissue, a metagene, and a cell-type node. For all metagenes *j* ∈ 1..*M*, we compute latent hyperedge attributes 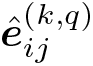 for a cell-type *q* in a tissue of interest *k* of individual *i* as follows:

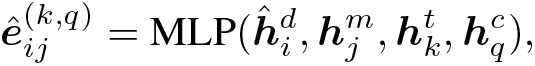

where 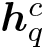 are parameters specific to each unique cell-type *q* and the MLP is shared for all combinations of metagenes, individuals, tissues, and cell-types.

##### Zero-inflated model

We employ the following probabilistic model:

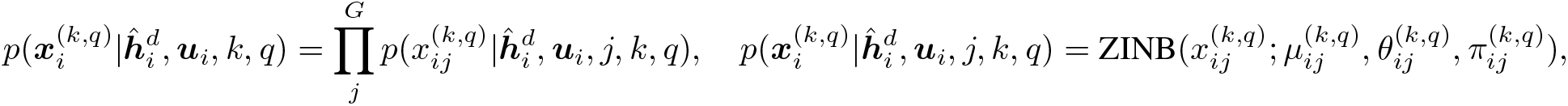

where ZINB is a zero-inflated negative binomial distribution. The mean 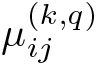, dispersion 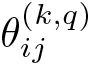, and dropout probability 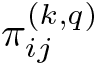 parameters are computed as:

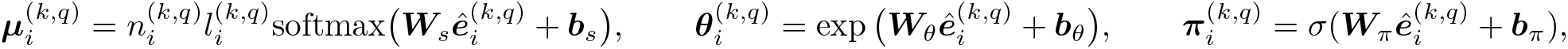

where ***W***_*s*_, ***W***_*θ*_, ***W***_*π*_, ***b***_*s*_, ***b***_*θ*_, and ***b***_*π*_ are learnable parameters; 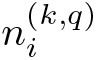 is the number of cells in the signature, and 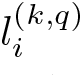 is their average library size. At train time, we set 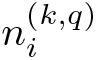 to match the ground truth number of cells. At test time, the number of cells 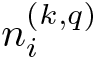 is user-definable. We model the average library size 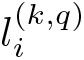 with a log-normal distribution

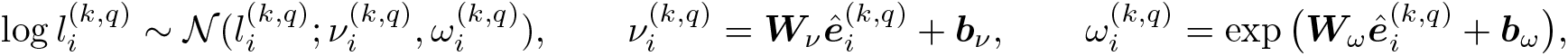

where ***W***_*ν*_, ***W***_*ω*_, ***b***_*ν*_, and ***b***_*ω*_ are learnable parameters. Optionally, we can use the observed library size.

##### Optimisation

Single-cell transcriptomic studies typically measure single-cell gene expression for a limited number of individuals, tissues, and cell-types, so aggregating single-cell profiles per individual, tissue, and cell-type often results in small sample sizes. To address this challenge, we apply transfer learning by pre-training the hypergraph model *f* on the multi-tissue imputation task (see Section 4.3.1) and then fine-tuning the parameters of the signature inference module (Section 4.3.2) on the cell-type signature profiles. Concretely, we minimise the loss:

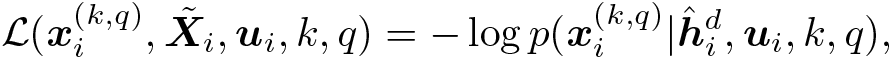

which corresponds to the negative log-likelihood of the observed cell-type signatures.

##### Inference of uncollected gene expression

To infer the signature of a cell-type *q* in a certain tissue *v* of interest, we first compute the latent individual embeddings 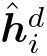 from the multi-tissue profiles 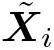 and then compute the mean of the distribution 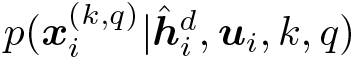 as 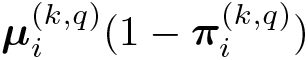. Alternatively, we can draw random samples from that distribution.

### 4.4 Data missingness assumption

By employing maximum likelihood inference on the observed data (Supplementary Materials), HYFA assumes that the data (i.e. tissues) are *Missing At Random* (MAR; Little and Rubin [2019]), that is, the missingness mechanism is independent of the unobserved data. Training HYFA via variational inference (Supplementary Materials) also necessitates the MAR assumption which, similar to Mattei and Frellsen [2019], arises from maximising the log-likelihood of the observed data through the Evidence Lower Bound (ELBO). The MAR assumption is less restrictive than the *Missing At Completely at Random* (MCAR) assumption — the missingness pattern is independent of the observed and unobserved data — of other methods such as mean imputation and GAIN [Yoon et al., 2018].

HYFA does not support data Missing Not At Random (MNAR), where the missingness mechanism depends on the unobserved data, i.e. the probability of being missing depends on unknown reasons [Van Buuren, 2018]. To handle this scenario, we would need to model the joint distribution 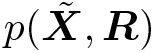 of the observed data 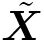 and the missingness mechanism **ℝ**, that is, the missingness mechanism would be nonignorable and would need to be explicitly modelled. This could be achieved through selection modeling [Heckman, 1976], which factorises the joint distribution as 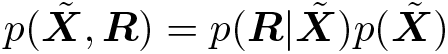, or pattern-mixture models [Glynn et al., 1986], which decompose the joint as 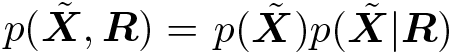. In general, it is impossible to test if MAR holds in a dataset [Schafer and Graham, 2002], but the impact of incorrectly assuming MAR is often minor [Collins et al., 2001].

### 4.5 Related work

#### Gene expression imputation methods

A standard approach for imputation of uncollected transcriptomics values is to use a proxy tissue (e.g. whole blood) as a surrogate [Hoon et al., 2000]. However, gene expression is known to be tissue and cell-type specific, limiting the effectiveness of this technique. Other related studies infer tissue-specific gene expression from genetic information. Wang et al. [2016] propose a mixed-effects model to infer uncollected data in multiple tissues from expression quantitative trait loci (eQTLs). Sul et al. [2013] introduce a model termed Meta-Tissue that aggregates information from multiple tissues to increase statistical power of eQTL detection. Nonetheless, these approaches do not model the complex relationships between measured and unmeasured gene expression traits among tissues and cell types, and individual-level genetic information (such as at eQTLs) is often unavailable and subject to privacy concerns. Instead, recent closely related work relies on linear factor analysis and dimensionality reduction techniques. TEEBoT (Tissue Expression Estimation using Blood Transcriptome) [Basu et al., 2021] projects gene expression from a single reference tissue (i.e. whole blood) into a low-dimensional space via principal component analysis (PCA), followed by linear regression to impute uncollected values. HYFA allows a departure from the linearity assumption of TEEBoT and also handles a variable number of reference tissues.

#### Knowledge graph embedding techniques

Our framework leverages ideas from knowledge graph embedding techniques by using learnable embeddings for biological entities (i.e. tissues, cell-types, and metagenes). Since the advent of word embeddings [Mikolov et al., 2013], several approaches have emerged to learn vector representations of entities and relations in knowledge graphs [Bordes et al., 2013, Wang et al., 2014, Trouillon et al., 2016, Dettmers et al., 2018]. TransE [Bordes et al., 2013] represents entities as low-dimensional vectors and relationships as translations in the embedding space, and optimises parameters through an energy-based objective. TransH [Wang et al., 2014] extends the TransE framework by projecting the entity embeddings into relation-specific hyperplanes. ComplEx [Trouillon et al., 2016] utilises complex vectors that capture antisymmetric entity relations. ConvE [Dettmers et al., 2018] models interactions between input entities and relations through convolutional and fully-connected layers. Despite all the recent advances, knowledge graph embeddings have been understudied for modelling higher-order structures (i.e. hyperedges). Moreover, while these methods are capable of link prediction, they are limited to a transductive setting, where the full set of entities (e.g. individuals) must be known at train time [Teru et al., 2020].

#### Graph representation learning

Graph neural networks remedy this problem by leveraging the structure and properties of graphs to compute node features, allowing to handle unseen entities at inference time (e.g. individuals). Graph neural networks operating on hypergraphs have recently started to flourish, with approaches such as HEAT [Georgiev et al., 2022] and *rxn-hypergraph* [Tavakoli et al., 2022] attaining state-of-the-art results in tasks involving higher-order relationships, such as source code [Georgiev et al., 2022] and chemical reactions [Tavakoli et al., 2022]. In terms of graph-based imputation methods, the closest approach to our framework is GRAPE [You et al., 2020]. GRAPE represents tabular data as a bipartite graph, where observations and features are two types of nodes, and the observed feature values are attributed edges between the nodes [You et al., 2020]. Imputation of missing features then corresponds to an edge-level prediction task. HYFA subsumes GRAPE’s bigraph in that our hypergraph becomes a bipartite graph when a single metagene is employed. This allows for a trade-off between feature granularity and over-squashing, which happens when information from a large receptive field is compressed into fixed-length node vectors [Alon and Yahav, 2020]. In terms of message passing, HYFA distinguishes between dynamic nodes (udpated during message passing) and static nodes (with learnable node features that are not updated during message passing), eliminating the dependency of tissue and metagene representations on donor features and, by transitivity, undesired dependencies between inviduals. HYFA is thus a hybrid and flexible approach that combines features from knowledge graph embedding and graph representation learning techniques.

#### Single-cell variational inference

Our framework is related to single-cell variational inference (scVI) [Lopez et al., 2018] in that it can also be optimised via variational inference (Supplementary Materials), e.g. via a (zero-inflated) negative binomial likelihood, treating the individual representations as latent variables. In contrast to scVI, however, HYFA offers features to handle a variable number of reference tissues. It also incorporates inductive biases to reuse knowledge across tissues, allowing the model to scale to larger multi-tissue samples.

### 4.6 eQTL mapping

The breadth of tissues in the GTEx (v8) collection enabled us to comprehensively evaluate the extent to which eQTL discovery could be improved through the HYFA-imputed transcriptome data. We mapped eQTLs that act in cis to the target gene (cis-eQTLs), using all SNPs within ± 1 Mb of the transcription start site of each gene. For the imputed and the original (incomplete) datasets, we considered SNPs significantly associated with gene expression, at a false discovery rate ≤ 0.10. We applied the same GTEx eQTL mapping pipeline, as previously described [Consortium, 2015], to the imputed and original datasets to quantify the gain in eQTL discovery from the HYFA-imputed dataset.

### 4.7 Pathway enrichment analysis

Similar to [Zhao et al., 2021], we employed Gene Set Enrichment Analysis (GSEA) [Subramanian et al., 2005] to relate HYFA’s metagene factors to known biological pathways. This is advantageous to over-representation analysis, which requires selecting an arbitrary cutoff to select enriched genes. GSEA, instead, computes a running sum of enrichment scores by descending a sorted gene list [Subramanian et al., 2005, Zhao et al., 2021].

We applied GSEA to the gene loadings in HYFA’s encoder. Specifically, let ***W***_*j*_ ∈ ℝ^*F* ×*G*^ be the gene loadings for metagene *j*, where *F* is the number of factors (i.e. number of hyperedge attributes) and *G* is the number of genes (Equation 1). For every factor in ***W***_*j*_, we employed `blitzGSEA` [Lachmann et al., 2022] to calculate the running sum of enrichment scores by descending the gene list sorted by the factor’s gene loadings. The enrichment score for a query gene set is the maximum difference between *p*_*hit*_(𝒮, *i*) and *p*_*miss*_(𝒮, *i*) [Zhao et al., 2021], where *p*_*hit*_(𝒮, *i*) is the proportion of genes in 𝒮 weighted by their gene loadings up to gene index *i* in the sorted list [Zhao et al., 2021]. We then calculated pathway enrichment p-values through a permutation test (with *N* = 100 trials) by randomly shuffling the gene list. We employed the Benjamini-Hochberg method to correct for multiple testing.

### 4.8 GTEx bulk and single-nucleus RNA-seq data processing

The GTEx dataset is a public resource that has generated a broad collection of gene expression data collected from a diverse set of human tissues [Consortium, 2020]. We downloaded the data from the GTEx portal (Data availability). After the processing step, the GTEx-v8 dataset consisted of 15197 samples (49 tissues, 834 donors) and 12557 genes. The dataset was randomly split into 500 train, 167 validation, and 167 test donors. Each donor had an average of 18.22 collected tissues. The processing steps are described below.

#### Normalised bulk transcriptomics (GTEx-v8)

Following the GTEx eQTL discovery pipeline (https://github.com/broadinstitute/gtex-pipeline/tree/master/qtl), we processed the data as follows:

1. Discard underrepresented tissues (n=5), namely bladder, cervix (ectocervix, endocervix), fallopian tube, and kidney (medulla).
2. Select set of overlapping genes across all tissues.
3. Discard donors with only one collected tissue (n=4).
4. Select genes based on expression thresholds of ≥ 0.1 transcripts per kilobase million (TPM) in ≥ 20% of samples and ≥ 6 reads (unnormalised) in ≥ 20% of samples.
5. Normalise read counts across samples using the trimmed mean of M values (TMM) method [Robinson and Oshlack, 2010].
6. Apply inverse normal transformation to the expression values for each gene.

#### Cell-type signatures from a paired snRNA-seq dataset (GTEx-v9)

We downloaded paired snRNA-seq data for 16 GTEx individuals [Eraslan et al., 2021] (Data availability) collected in 8 GTEx tissues, namely skeletal muscle, breast, esophagus (mucosa, muscularis), heart, lung, prostate, and skin. We split the 16 individuals into train, validation, and test donors according to the GTEx-v8 split. We processed the data as follows:

1. Select set of overlapping genes between bulk RNA-seq (GTEx-v9) and paired snRNA-seq dataset [Eraslan et al., 2021].
2. Select top 3000 variable genes using the Scanpy function `scanpy.pp.highly_variable_genes` with flavour setting `seurat_v3` [Wolf et al., 2018, Stuart et al., 2019].
3. Discard underrepresented cell-types occurring in less that 10 tissue-individual combinations.
4. Aggregate (i.e. sum) read counts by individual, tissue, and (broad) cell-type. This resulted in a dataset of 226 unique signatures.

### 4.9 Implementation and reproducibility

We report the selected hyperparameters in Supplementary Materials H. HYFA is implemented in Python [Van Rossum and Drake Jr, 1995]. Our framework and implementation are flexible (i.e. we support k-uniform hypergraphs), may be integrated in other bioinformatics pipelines, and may be useful for other applications in different domains. We used PyTorch [Paszke et al., 2019] to implement the model and scanpy [Wolf et al., 2018] to process the gene expression data. We performed hyperparameter optimisation with wandb [Biewald, 2020]. We employed blitzGSEA [Lachmann et al., 2022] for the pathway enrichment analysis. We also used NumPy [Harris et al., 2020], scikit-learn [Pedregosa et al., 2011], pandas [Wes McKinney, 2010], matplotlib [Hunter, 2007], and seaborn [Waskom, 2021]. Figure 1 was created with BioRender.com.

## Code availability

HYFA is publicly available at https://github.com/rvinas/HYFA.

## Data availability

The datasets analysed for this study, including bulk RNA-seq Consortium [2020] and snRNA-seq Eraslan et al. [2021], can be found in the GTEx portal: https://gtexportal.org/. A detailed summary of the GTEx samples and donor information can be found at: https://gtexportal.org/home/tissueSummaryPage. The full catalog of HYFA-derived eQTLs is downloadable at: https://zenodo.org/deposit/6815784.

## Acknowledgements

We thank Pietro Barbiero, David Buterez, Iulia Duta, Janine Lux, Andrei Margeloiu, Jacob Moss, Paul Scherer, and Nikola Simidjievski for useful feedback and discussions. The project leading to these results has received funding from Fundación Rafael del Pino (RV), with support from the Engineering and Physical Sciences Research Council (EPSRC DTG). PL was supported by MICA: Mental Health Data Pathfinder: University of Cambridge, Cambridgeshire and Peterborough NHS Foundation Trust, Microsoft, and the Medical Research Council (MC PC 17213). ERG acknowledges support from the following National Institutes of Health (NIH) grants: Genomic Innovator Award R35HG010718, NHGRI R01HG011138, NIMH R01MH126459, and NIA AG068026. We thank Phillip Lin for technical assistance and Vanderbilt’s Advanced Computing Center for Research and Education (ACCRE) for infrastructure support. We thank Janine Lux for support with the figures.

## A. Prediction scores for different accessible tissues as reference

**Extended Data Figure 1:**
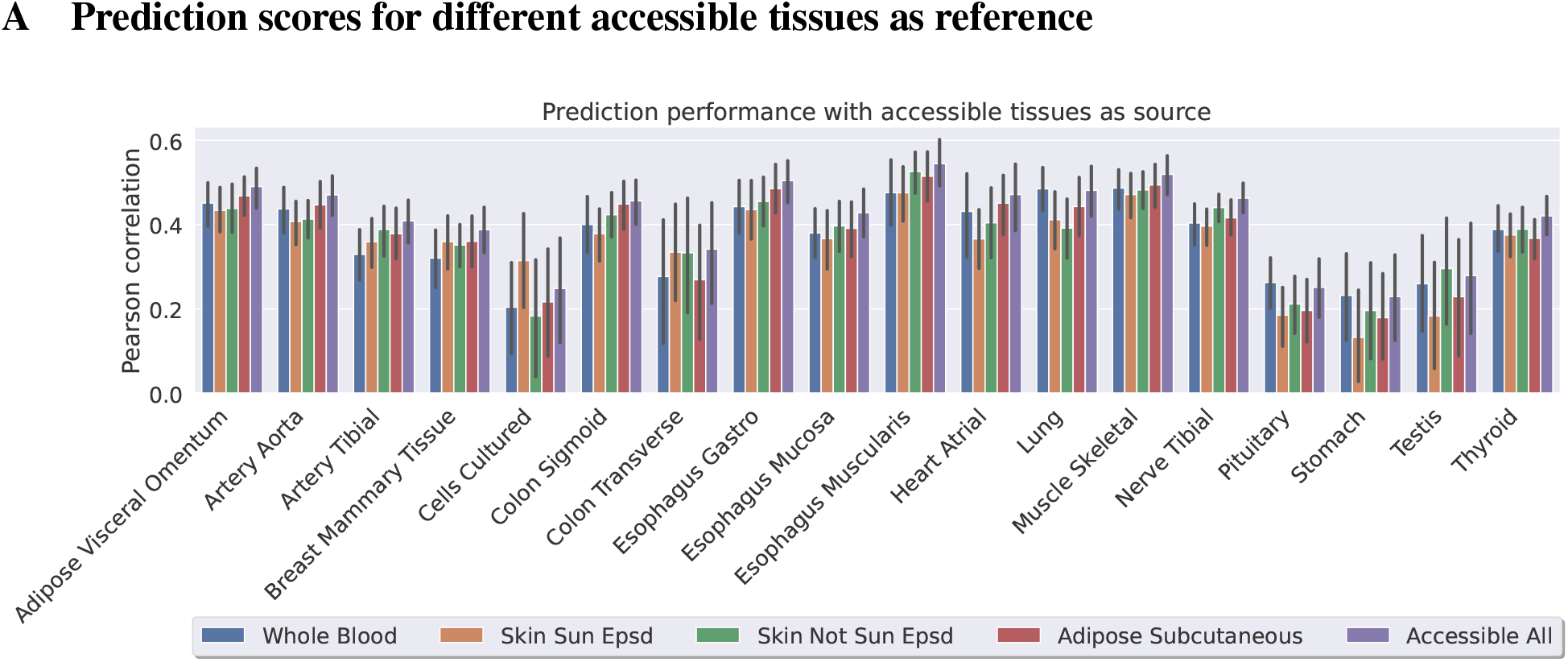
Prediction scores for different accessible tissues as reference. For each target tissue, we predicted the expression values based on accessible tissues (whole blood, skin sun exposed, skin not sun exposed, and adipose subcutaneous). We report the Pearson correlation coefficient between the predicted values and the actual gene expression values. For any given target tissue, we used the same set of individuals to evaluate performance, namely individuals in the test set with collected gene expression measurements in all the corresponding tissues. Target tissues represented by less than 25 test individuals were discarded. Error bars indicate 95% confidence intervals. The method attains the best performance in 14 out of 18 tissues when all accessible tissues are simultaneously used as reference.

## B Train-test splits of cell-type signatures

**Extended Data Figure 2:**
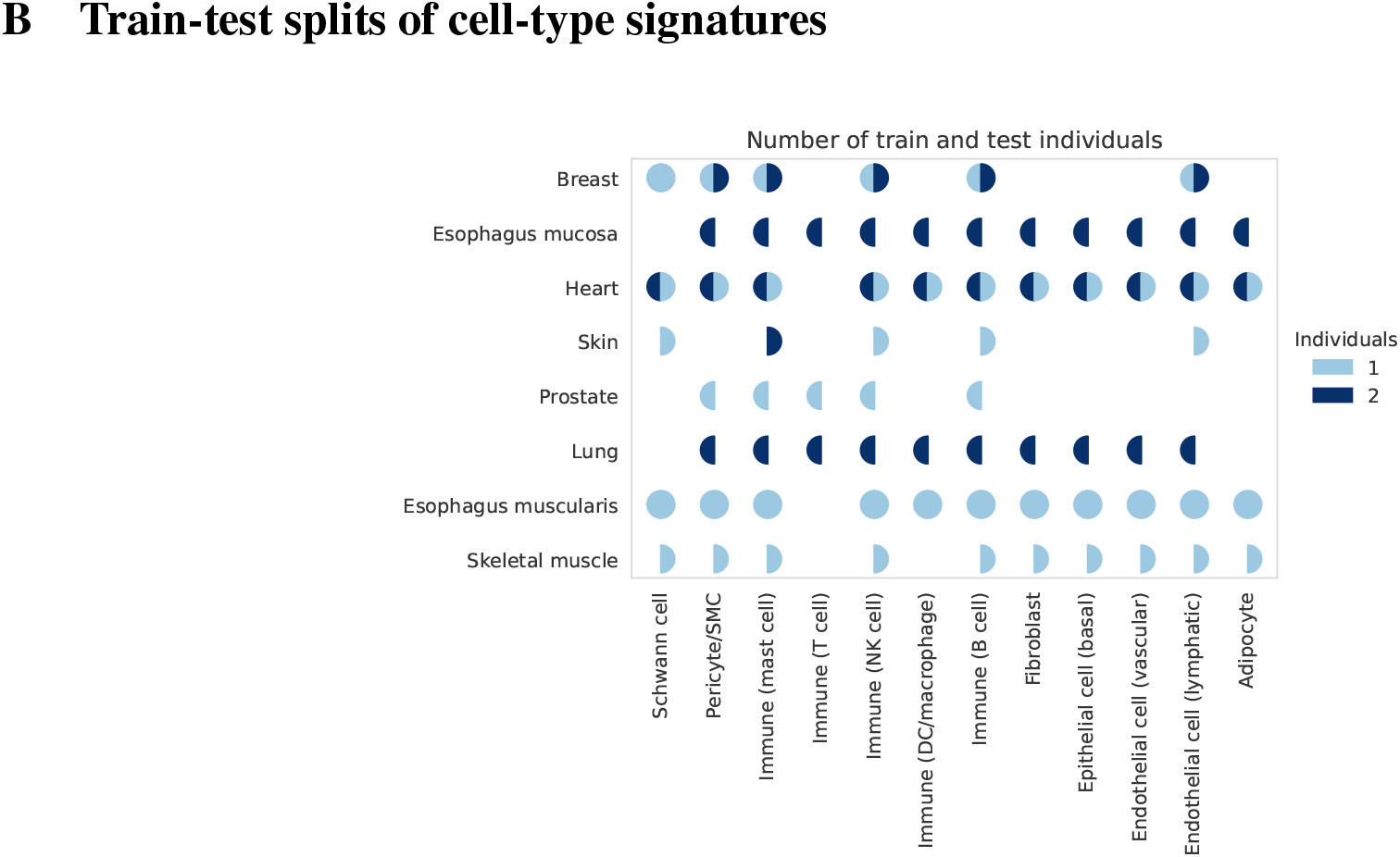
Number of train (left semi-circle) and test (right semi-circle) signatures per tissue and cell-type. Each signature corresponds to the aggregated tissue- and cell-type-specific scRNA-seq counts for a given individual. For any combination of tissue and cell-type, there are no more than 2 individual-specific signatures in the same set. Blank semi-circles indicate zero signatures. Note that some signatures (e.g. cell-types in skeletal muscle) are only present in the test set.

## C Prediction of cell-type signatures with inferred library sizes

**Extended Data Figure 3:**
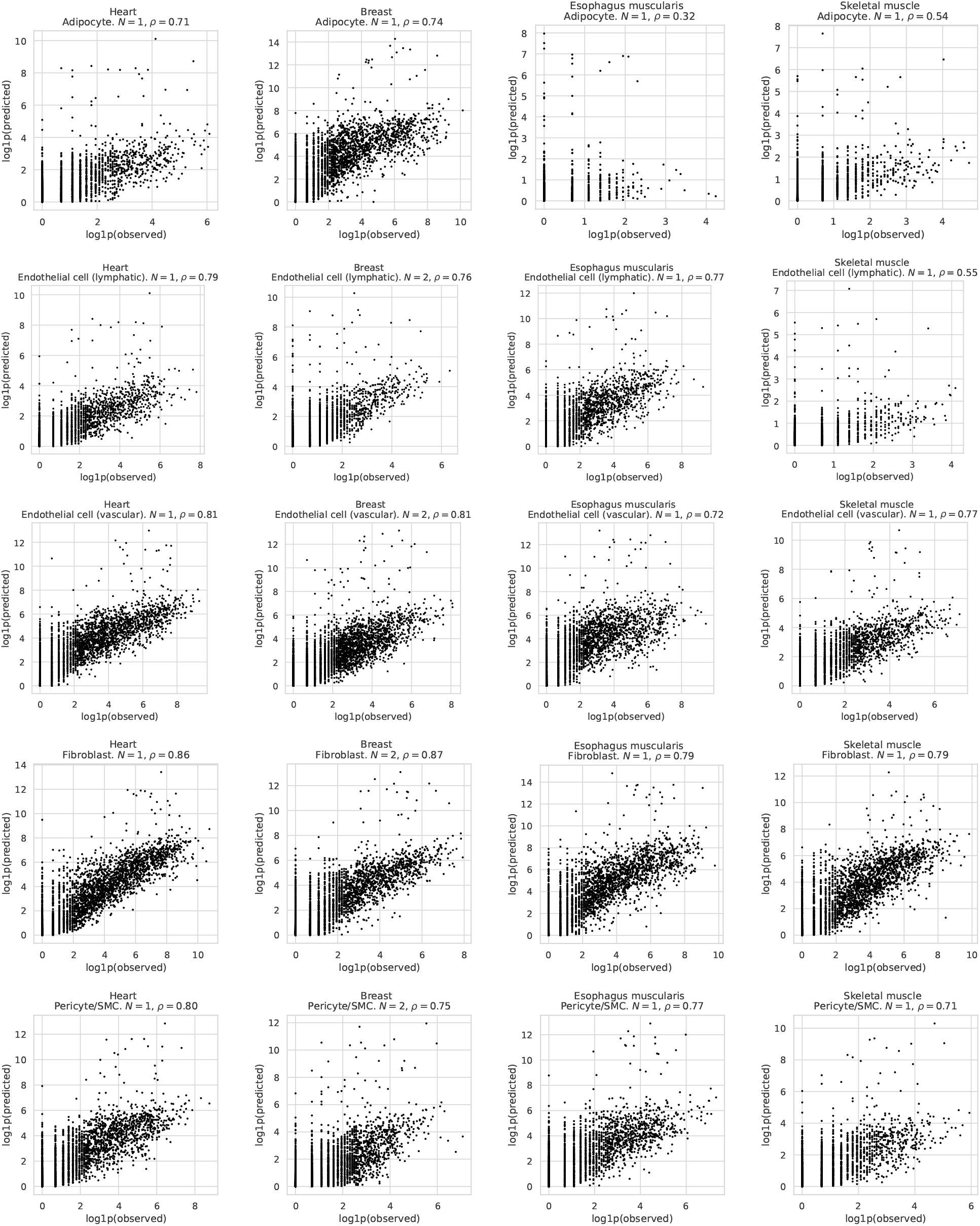
Prediction of cell-type signatures. HYFA imputes individual- and tissue-specific cell-type signatures from bulk multi-tissue gene expression. The scatter plots depict the Pearson correlation *ρ* between the logarithmised ground truth and predicted signatures for *N* unseen individuals. To predict the signatures, we inferred the library sizes 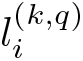 and used the observed number of cells 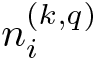.

## D GTEx statistics

### D.1 Histogram with samples per tissue

**Extended Data Figure 4:**
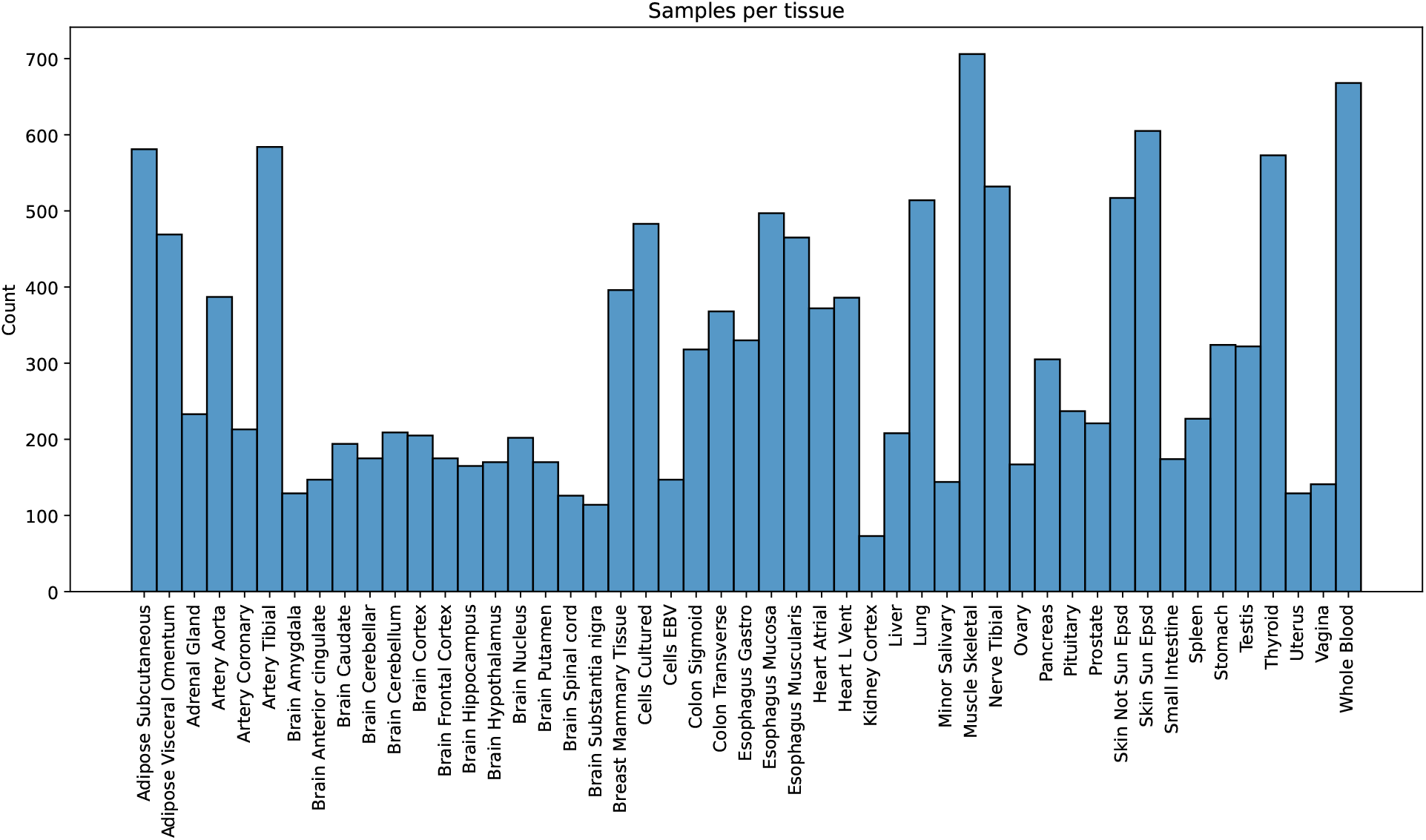
Number of samples per tissue

### D.2 Small overlap between brain and gastrointestinal tissues

**Extended Data Figure 5:**
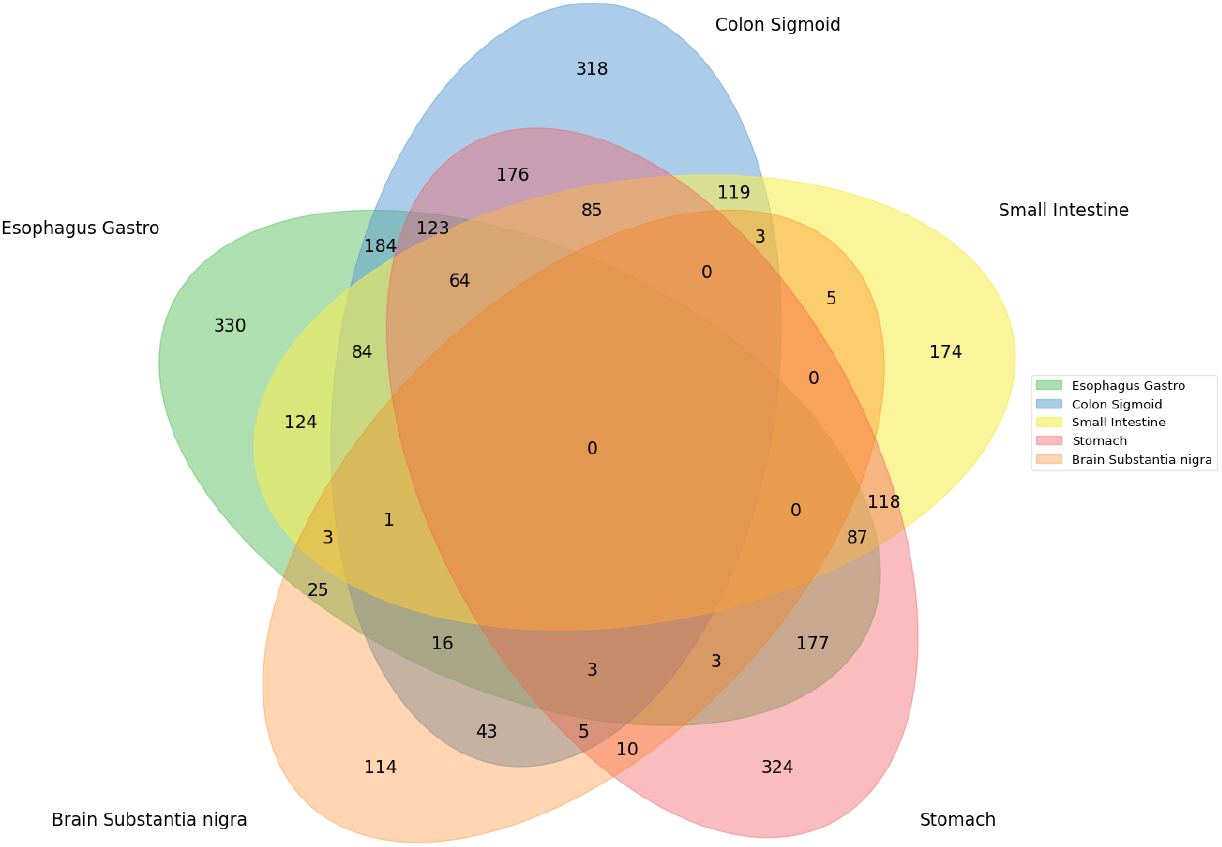
Overlap between brain and gastrointestinal tissues in terms of GTEx individuals.

## E. Connection with maximum likelihood

Let x_*obs*_ be a random variable denoting the observed data (e.g. multi-tissue gene expression with missing values corresponding to uncollected tissues). Our optimisation procedure (Methods) splits x_*obs*_ into *pseudo-observed* 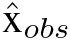 and *pseudo-missing* 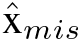 values, that is, 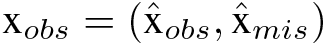. The log-likelihood of the observed data then corresponds to:

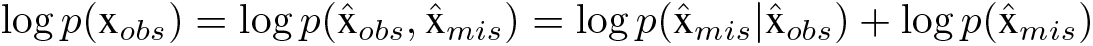

and 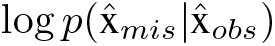 is precisely the quantity that our loss function is maximising through the pseudo-mask mechanism (Methods).

## F Training algorithm

### Optimisation

We optimise the model to minimise the mean squared error *L* between the normalised, ground-truth gene expression 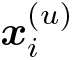 and the imputed values 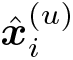:

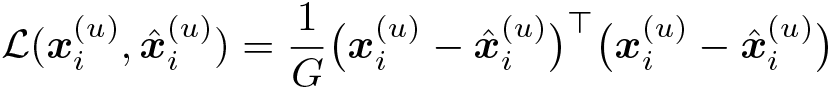

where *G* is the number of genes. At train time, for any given individual, we dynamically mask out the expression values of a measured tissue type at random and treat them as uncollected, i.e. the ground truth. Algorithm 1 summarises the training algorithm.

#### Algorithm 1 Training algorithm

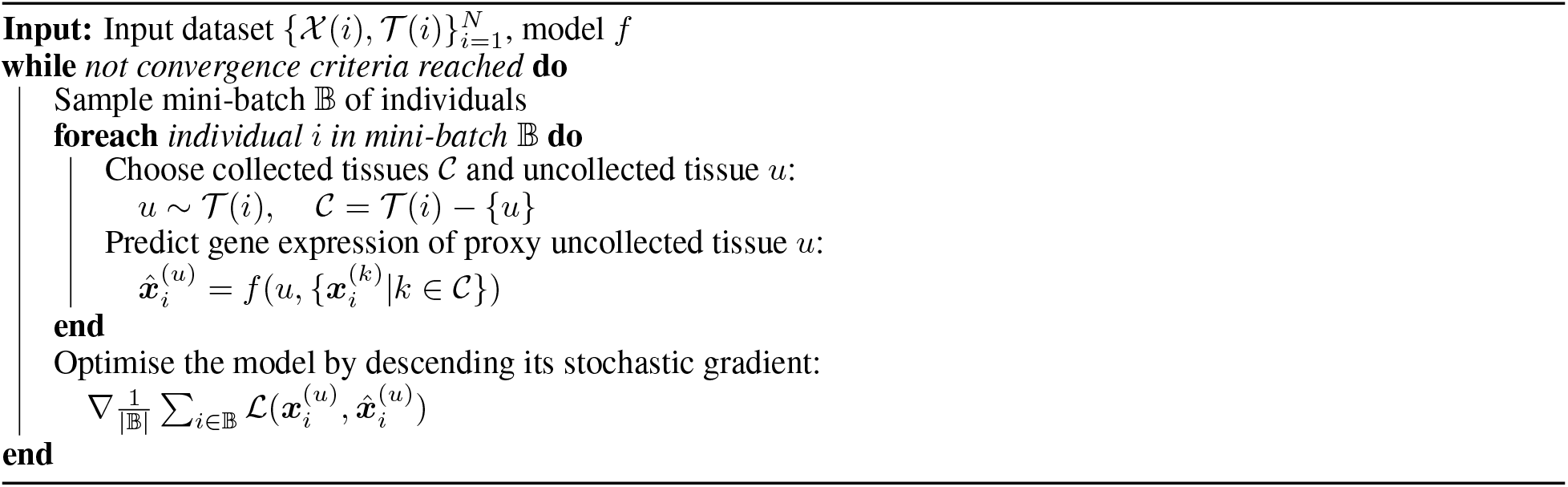

## G Training HYFA via variational inference

HYFA can alternatively be trained via variational inference by introducing a variational distribution 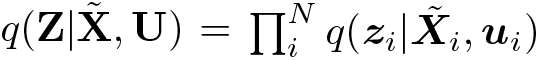, where ***z***_*i*_ is a latent variable that explains the high-dimensional, multi-tissue gene expression data.

### Parameters of inference model

Given the updated donor representations 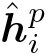, we compute the parameters of the inference model 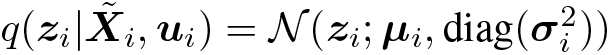 as follows:

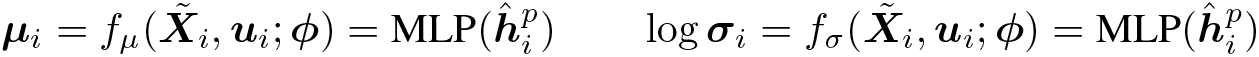

where MLP denotes a multilayer perceptron.

### Parameters of generative model

Assuming a Gaussian likelihood, for a given sample 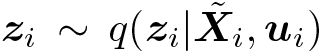, we compute the parameters of the generative model 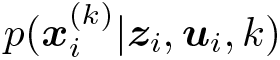 as follows:

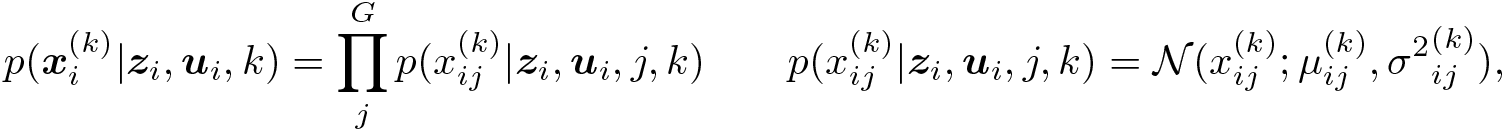

where the mean 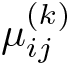 and standard deviation 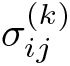 are computed as follows:

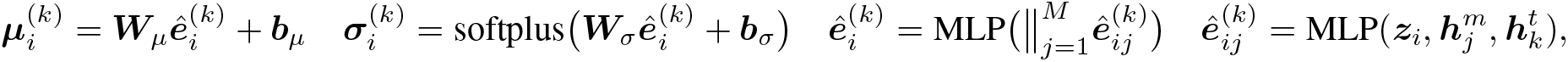

### Optimisation

We maximise the evidence lower bound on the data log-likelihood:

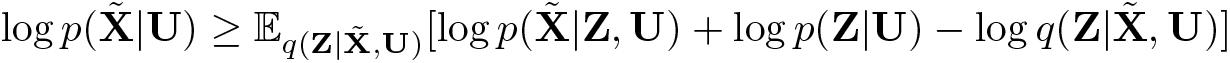

where the prior *p*(**Z**|**U**) is a factorised normal distribution conditioned on demographic information:

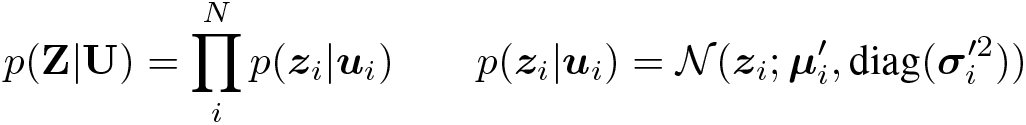

with parameters 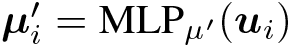 and log 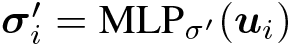. Importantly, leveraging a factorised prior conditioned on auxiliary variables guarantees identifiability under certain conditions [Khemakhem et al., 2020].

### Inference of uncollected gene expression measurements

We infer the gene expression values 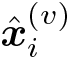 of an uncollected tissue *v* from a given donor *i* as follows:

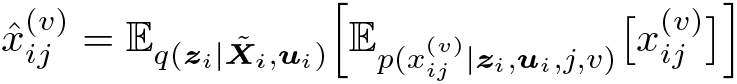

In other words, given the multi-tissue gene expression 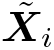 and demographic information ***u***_*i*_, we compute the expectation of the target gene expression 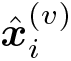 over the inference and generative models.

## H Ablation of architecture

We ablate the impact of two key architectural components of HYFA: (1) representating multi-tissue gene expression as a hypergraph of individuals, metagenes, and tissues; and (2) the design of a specialised hypergraph message passing neural network layer with attentional aggregation.

### Number of metagenes

In Figure 6, we plot the validation loss and correlation coefficient vs. the number of metagenes for both attentional (GAT) and standard message passing (MPNN). The attentional model (GAT) refers to HYFA with an attention-based aggregation mechanism (Methods), while the message passing model (MPNN) refers to HYFA with simple average aggregation (i.e. mean across all incoming messages). For each number of metagenes, we ran hyperparameter optimisation with wandb [Biewald, 2020] to obtain the loss and Pearson correlation coefficient *ρ* for the best performing model (we ran sweeps with a maximum of 100 runs). For fair comparison across runs, validation metrics were computed for a fixed subset of target tissues: ‘Lung’, ‘Pancreas’, ‘Heart_Atrial’, and ‘Esophagus_Muscularis’. The hyperparameter values considered for ablation studies are available in Extended Data Table 2.

As noted in Section 4.2.1, modulating the number of metagenes controls the growth of the receptive field for each node in the hypergraph and helps alleviate over-squashing. Setting very low number of metagenes is computationally fast during training and inference but may compress fine-grained information (e.g. setting metagenes to 1 results in a bipartite graph of individuals and tissues), while a very high number of metagenes preserves fine-grained relationships between genes, tissues, and individuals but may become computationally intractable. Figure 6 shows that there is a ‘sweet spot’ for the number of metagenes between 50-100. Using over 200 metagenes can consume upwards of 32 GB of GPU memory with very long iteration times, and we subsequently run into ‘out-of-memory’ errors.

### Hypergraph message passing architecture

Extended Data Table 1 summarises results for the best performing GAT and MPNN, demonstrating that the specialised hypergraph attentional aggregation brings notable gains in imputation performance. This is consistent with the observation that, through the attention mechanism, the model can prioritise certain messages from others, alleviating the over-squashing problem. As a naïve baseline, we also show results for a structure-agnostic model which does not perform any message passing and simply predicts the hyperedge attributes via an MLP, as in Section 4.3.1. The inductive bias of reusing knowledge across tissues via message passing seems critical for gene expression imputation.

**Extended Data Figure 6:**
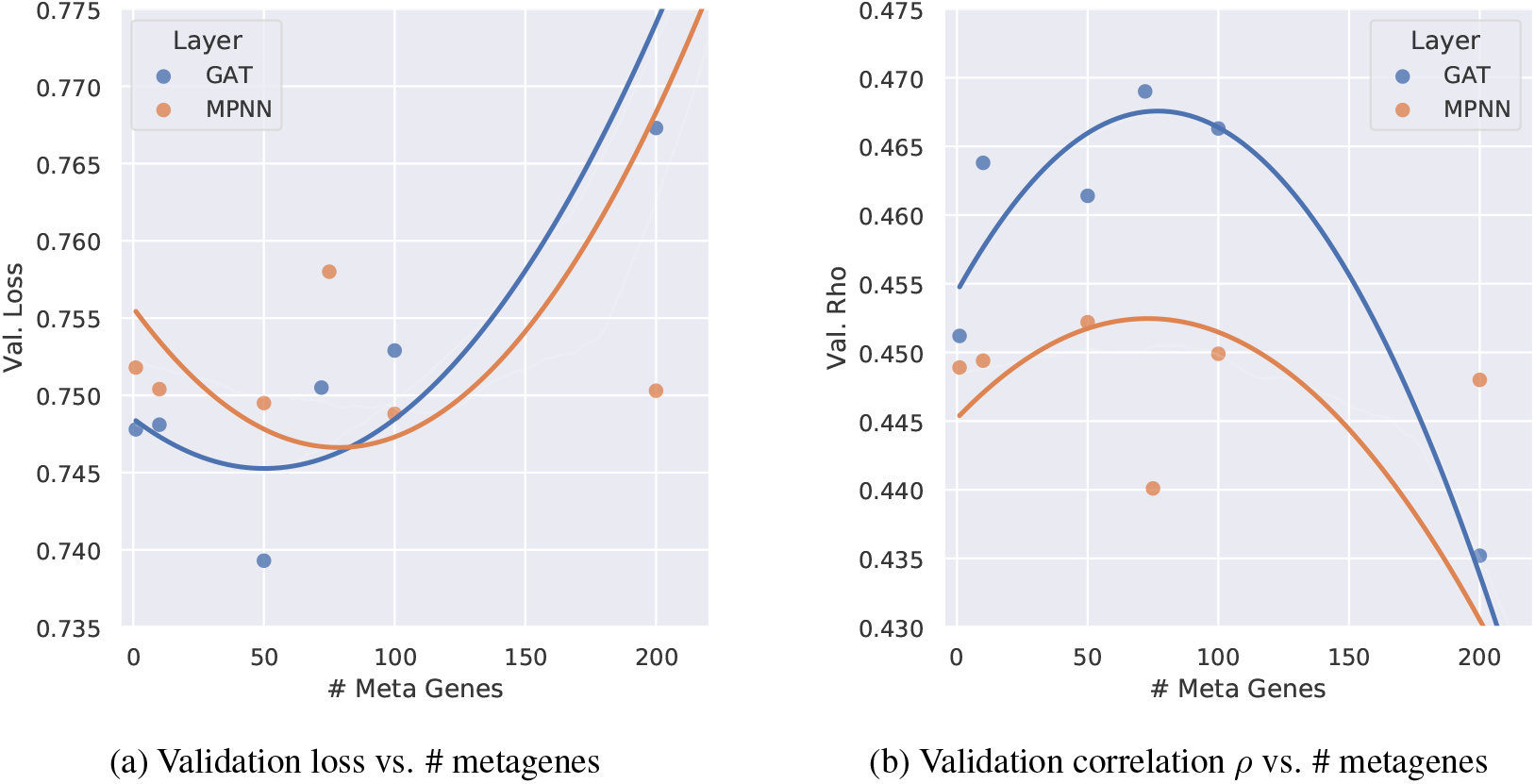
Impact of number of metagenes in hypergraph representations vs. model performance. There is a ‘sweet spot’ for the number of metagenes between 50-100 that leads to optimal performance for both attentional (GAT) and standard message passing (MPNN). Curves are estimated via a polynomial regression with order 2.

**Extended Data Table 1:** Ablation study of hypergraph message passing design. A specialised hypergraph attentional aggregation brings significant gains in imputation performance over standard message passing as well as a naïve structure-agnostic baseline.

| GNN Layer | Val. Loss ↓ | Val. Correlation ↑ |
| --- | --- | --- |
| Structure-agnostic MLP | 0.9719 | 0.0396 |
| Message Passing (MPNN) | 0.7488 | 0.4499 |
| Attentional (GAT) | **0.7393** | **0.4614** |

**Extended Data Table 2:** Hyperparamter values considered for ablation studies. We used wandb [Biewald, 2020] to run Bayesian hyperparameter search over the variables and ranges considered. Note that with the Attentional GAT layer, the total dimension of the message ***m***_*ijk*_ is multiplied by the number of attention heads (here, 28 × 28 = 784).

| Hyperparameter | Values Considered | Best Value |
| --- | --- | --- |
| GNN Layer | { GAT, MPNN } | GAT |
| Num. Metagenes | 10 – 200 | 50 |
| Num. Message Passing Layers | 1 – 3 | 2 |
| Num. MLP Layers (within GNN) | 1 – 2 | 1 |
| Num. MLP Layers (Prediction head) | 1 – 2 | 2 |
| Num. Attention Heads (GAT only) | 4 – 32 | 28 |
| Dimension of Donor Emb. $h^d$ | 16 – 128 | 71 |
| Dimension of Metagene Emb. $h^m$ | 16 – 128 | 48 |
| Dimension of Tissue Emb. $h^t$ | 16 – 128 | 120 |
| Dimension of Hyperedge Attr. $e_{ij}$ | 16 – 128 | 98 |
| Dimension of Message $m_{ijk}$ | 16 – 128 | 28 |
| Learning Rate | 0.0001 – 0.005 | 0.00045 |
| Batch Size | 16 – 64 | 63 |
| Dropout | 0.0 – 0.5 | 0.17385 |
| Normalisation | { BatchNorm, LayerNorm, None } | BatchNorm |
| Activation Function | { ReLU, Swish } | Swish |

## References

Mahashweta Basu, Kun Wang, Eytan Ruppin, and Sridhar Hannenhalli. Predicting tissue-specific gene expression from whole blood transcriptome. Science Advances, 7(14):eabd6991, 2021.

GTEx Consortium. The gtex consortium atlas of genetic regulatory effects across human tissues. Science, 369(6509): 1318–1330, 2020.

Xiaonan Yang, Ling Kui, Min Tang, Dawei Li, Kunhua Wei, Wei Chen, Jianhua Miao, and Yang Dong. High-throughput transcriptome profiling in drug and biomarker discovery. Frontiers in genetics, 11:19, 2020.

Chenling Xu, Romain Lopez, Edouard Mehlman, Jeffrey Regier, Michael I Jordan, and Nir Yosef. Probabilistic harmonization and annotation of single-cell transcriptomics data with deep generative models. Molecular systems biology, 17(1):e9620, 2021.

Dave SB Hoon, Peter Bostick, Christine Kuo, Tetsuro Okamoto, He-Jing Wang, Robert Elashoff, and Donald L Morton. Molecular markers in blood as surrogate prognostic indicators of melanoma recurrence. Cancer research, 60(8): 2253–2257, 2000.

Chaochao Cai, Peter Langfelder, Tova F Fuller, Michael C Oldham, Rui Luo, Leonard H van den Berg, Roel A Ophoff, and Steve Horvath. Is human blood a good surrogate for brain tissue in transcriptional studies? BMC genomics, 11 (1):1–15, 2010.

Geoffrey Istas, Ken Declerck, Maria Pudenz, Veronica Lendinez-Tortajada, Montserrat Leon-Latre, Karen Heyninck, Guy Haegeman, Jose A Casasnovas, Maria Tellez-Plaza, Clarissa Gerhauser, et al. Identification of differentially methylated brca1 and crisp2 dna regions as blood surrogate markers for cardiovascular disease. Scientific reports, 7 (1):1–14, 2017.

Eric R Gamazon, Ayellet V Segrè, Martijn Van De Bunt, Xiaoquan Wen, Hualin S Xi, Farhad Hormozdiari, Halit Ongen, Anuar Konkashbaev, Eske M Derks, François Aguet, et al. Using an atlas of gene regulation across 44 human tissues to inform complex disease-and trait-associated variation. Nature genetics, 50(7):956–967, 2018.

Kayoung Kim, Min-Ji Kim, Da Won Kim, Su Yeong Kim, Steve Park, and Chan Beum Park. Clinically accurate diagnosis of alzheimer’s disease via multiplexed sensing of core biomarkers in human plasma. Nature communications, 11(1):1–9, 2020.

Dan Zhou, Yi Jiang, Xue Zhong, Nancy J Cox, Chunyu Liu, and Eric R Gamazon. A unified framework for joint-tissue transcriptome-wide association and mendelian randomization analysis. Nature genetics, 52(11):1239–1246, 2020.

Jiebiao Wang, Eric R Gamazon, Brandon L Pierce, Barbara E Stranger, Hae Kyung Im, Robert D Gibbons, Nancy J Cox, Dan L Nicolae, and Lin S Chen. Imputing gene expression in uncollected tissues within and beyond gtex. The American Journal of Human Genetics, 98(4):697–708, 2016.

Jae Hoon Sul, Buhm Han, Chun Ye, Ted Choi, and Eleazar Eskin. Effectively identifying eqtls from multiple tissues by combining mixed model and meta-analytic approaches. PLoS genetics, 9(6):e1003491, 2013.

Gokcen Eraslan, Eugene Drokhlyansky, Shankara Anand, Ayshwarya Subramanian, Evgenij Fiskin, Michal Slyper, Jiali Wang, Nicholas Van Wittenberghe, John M. Rouhana, Julia Waldman, Orr Ashenberg, Danielle Dionne, Thet Su Win, Michael S. Cuoco, Olena Kuksenko, Philip A. Branton, Jamie L. Marshall, Anna Greka, Gad Getz, Ayellet V. Segrè, François Aguet, Orit Rozenblatt-Rosen, Kristin G. Ardlie, and Aviv Regev. Single-nucleus cross-tissue molecular reference maps to decipher disease gene function. bioRxiv, 2021. doi:10.1101/2021.07.19.452954. URL https://www.biorxiv.org/content/early/2021/07/19/2021.07.19.452954.

Jean-Philippe Brunet, Pablo Tamayo, Todd R Golub, and Jill P Mesirov. Metagenes and molecular pattern discovery using matrix factorization. Proceedings of the national academy of sciences, 101(12):4164–4169, 2004.

Soumya Raychaudhuri, Joshua M Stuart, and Russ B Altman. Principal components analysis to summarize microarray experiments: application to sporulation time series. In Biocomputing 2000, pages 455–466. World Scientific, 1999.

Valentine Svensson, Adam Gayoso, Nir Yosef, and Lior Pachter. Interpretable factor models of single-cell RNA-seq via variational autoencoders. Bioinformatics, 36(11):3418–3421, 03 2020. ISSN 1367-4803. doi:10.1093/bioinformatics/btaa169. URL https://doi.org/10.1093/bioinformatics/btaa169.

Justin Gilmer, Samuel S Schoenholz, Patrick F Riley, Oriol Vinyals, and George E Dahl. Neural message passing for quantum chemistry. In International conference on machine learning, pages 1263–1272. PMLR, 2017.

Till Roenneberg and Martha Merrow. The circadian clock and human health. Current biology, 26(10):R432–R443, 2016.

Jean-Michel Davière and Patrick Achard. Organ communication: Cytokinins on the move. Nature plants, 3(8):1–2, 2017.

Sue C Bodine, Heddwen L Brooks, Nigel W Bunnett, Hilary A Coller, Mark R Frey, Bina Joe, Thomas R Kleyman, Merry L Lindsey, Andre Marette, Rory E Morty, et al. An american physiological society cross-journal call for papers on “inter-organ communication in homeostasis and disease”, 2021.

Leland McInnes, John Healy, and James Melville. Umap: Uniform manifold approximation and projection for dimension reduction. arXiv preprint arXiv:1802.03426, 2018.

Laurens Van der Maaten and Geoffrey Hinton. Visualizing data using t-sne. Journal of machine learning research, 9 (11), 2008.

Sandip Ray, Markus Britschgi, Charles Herbert, Yoshiko Takeda-Uchimura, Adam Boxer, Kaj Blennow, Leah Friedman, Douglas Galasko, Marek Jutel, Anna Karydas, Jeffrey Kaye, Jerzy Leszek, Bruce Miller, Lennart Minthon, Joseph Quinn, Gil Rabinovici, William Robinson, Marwan Sabbagh, Yuen So, and Tony Wyss-Coray. Classification and prediction of clinical alzheimer’s diagnosis based on plasma signaling proteins. Nature Medicine, 13:1359–1362, 10 2007. doi:10.1038/nm1653.

Kasper Lage, Niclas Tue Hansen, E Olof Karlberg, Aron C Eklund, Francisco S Roque, Patricia K Donahoe, Zoltan Szallasi, Thomas Skøt Jensen, and Søren Brunak. A large-scale analysis of tissue-specific pathology and gene expression of human disease genes and complexes. Proceedings of the National Academy of Sciences, 105(52): 20870–20875, 2008.

F Alexander Wolf, Philipp Angerer, and Fabian J Theis. Scanpy: large-scale single-cell gene expression data analysis. Genome biology, 19(1):1–5, 2018.

Tim Stuart, Andrew Butler, Paul Hoffman, Christoph Hafemeister, Efthymia Papalexi, William M Mauck III, Yuhan Hao, Marlon Stoeckius, Peter Smibert, and Rahul Satija. Comprehensive integration of single-cell data. Cell, 177(7): 1888–1902, 2019.

Alexandra C Nica and Emmanouil T Dermitzakis. Expression quantitative trait loci: present and future. Philosophical Transactions of the Royal Society B: Biological Sciences, 368(1620):20120362, 2013.

Matthew V Rockman and Leonid Kruglyak. Genetics of global gene expression. Nature Reviews Genetics, 7(11): 862–872, 2006.

Maxim V Kuleshov, Matthew R Jones, Andrew D Rouillard, Nicolas F Fernandez, Qiaonan Duan, Zichen Wang, Simon Koplev, Sherry L Jenkins, Kathleen M Jagodnik, Alexander Lachmann, et al. Enrichr: a comprehensive gene set enrichment analysis web server 2016 update. Nucleic acids research, 44(W1):W90–W97, 2016.

Clair R Martin, Vadim Osadchiy, Amir Kalani, and Emeran A Mayer. The brain-gut-microbiome axis. Cellular and molecular gastroenterology and hepatology, 6(2):133–148, 2018.

Samuel Davis, Thomas H Aldrich, David M Valenzuela, Vivien Wong, Mark E Furth, Stephen P Squinto, and George D Yancopoulos. The receptor for ciliary neurotrophic factor. Science, 253(5015):59–63, 1991.

Sumei Liu. Neurotrophic factors in enteric physiology and pathophysiology. Neurogastroenterology & Motility, 30(10): e13446, 2018.

Baoji Xu and Xiangyang Xie. Neurotrophic factor control of satiety and body weight. Nature Reviews Neuroscience, 17(5):282–292, 2016.

Louis J Sparvero, Denise Asafu-Adjei, Rui Kang, Daolin Tang, Neilay Amin, Jaehyun Im, Ronnye Rutledge, Brenda Lin, Andrew A Amoscato, Herbert J Zeh, et al. Rage (receptor for advanced glycation endproducts), rage ligands, and their role in cancer and inflammation. Journal of translational medicine, 7(1):1–21, 2009.

Aravind Subramanian, Pablo Tamayo, Vamsi K Mootha, Sayan Mukherjee, Benjamin L Ebert, Michael A Gillette, Amanda Paulovich, Scott L Pomeroy, Todd R Golub, Eric S Lander, et al. Gene set enrichment analysis: a knowledge-based approach for interpreting genome-wide expression profiles. Proceedings of the National Academy of Sciences, 102(43):15545–15550, 2005.

Yifan Zhao, Huiyu Cai, Zuobai Zhang, Jian Tang, and Yue Li. Learning interpretable cellular and gene signature embeddings from single-cell transcriptomic data. Nature communications, 12(1):1–15, 2021.

Dylan Kotliar, Adrian Veres, M Aurel Nagy, Shervin Tabrizi, Eran Hodis, Douglas A Melton, and Pardis C Sabeti. Identifying gene expression programs of cell-type identity and cellular activity with single-cell rna-seq. Elife, 8, 2019.

Jiaxuan You, Xiaobai Ma, Daisy Ding, Mykel Kochenderfer, and Jure Leskovec. Handling missing data with graph representation learning. NeurIPS, 2020.

Antoine Bordes, Nicolas Usunier, Alberto Garcia-Duran, Jason Weston, and Oksana Yakhnenko. Translating embeddings for modeling multi-relational data. Advances in neural information processing systems, 26, 2013.

Uri Alon and Eran Yahav. On the bottleneck of graph neural networks and its practical implications. arXiv preprint arXiv:2006.05205, 2020.

Shaked Brody, Uri Alon, and Eran Yahav. How attentive are graph attention networks?, 2022.

Petar Veličković, Guillem Cucurull, Arantxa Casanova, Adriana Romero, Pietro Liò, and Yoshua Bengio. Graph attention networks, 2018.

Ashish Vaswani, Noam Shazeer, Niki Parmar, Jakob Uszkoreit, Llion Jones, Aidan N Gomez, L ukasz Kaiser, and Illia Polosukhin. Attention is all you need. In I. Guyon, U. V. Luxburg, S. Bengio, H. Wallach, R. Fergus, S. Vishwanathan, and R. Garnett, editors, Advances in Neural Information Processing Systems, volume 30. Curran Associates, Inc., 2017. URL https://proceedings.neurips.cc/paper/2017/file/3f5ee243547dee91fbd053c1c4a845aa-Paper.pdf.

Ramon Viñas, Tiago Azevedo, Eric R. Gamazon, and Pietro Liò. Deep learning enables fast and accurate imputation of gene expression. Frontiers in Genetics, 12, 2021. ISSN 1664-8021. doi:10.3389/fgene.2021.624128. URL https://www.frontiersin.org/article/10.3389/fgene.2021.624128.

Roderick JA Little and Donald B Rubin. Statistical analysis with missing data, volume 793. John Wiley & Sons, 2019.

Pierre-Alexandre Mattei and Jes Frellsen. Miwae: Deep generative modelling and imputation of incomplete data sets. In International conference on machine learning, pages 4413–4423. PMLR, 2019.

Jinsung Yoon, James Jordon, and Mihaela Schaar. Gain: Missing data imputation using generative adversarial nets. In International conference on machine learning, pages 5689–5698. PMLR, 2018.

Stef Van Buuren. Flexible imputation of missing data. CRC press, 2018.

James J Heckman. The common structure of statistical models of truncation, sample selection and limited dependent variables and a simple estimator for such models. In Annals of economic and social measurement, volume 5, number 4, pages 475–492. NBER, 1976.

Robert J Glynn, Nan M Laird, and Donald B Rubin. Selection modeling versus mixture modeling with nonignorable nonresponse. In Drawing inferences from self-selected samples, pages 115–142. Springer, 1986.

Joseph L Schafer and John W Graham. Missing data: our view of the state of the art. Psychological methods, 7(2):147, 2002.

Linda M Collins, Joseph L Schafer, and Chi-Ming Kam. A comparison of inclusive and restrictive strategies in modern missing data procedures. Psychological methods, 6(4):330, 2001.

Tomas Mikolov, Kai Chen, Greg Corrado, and Jeffrey Dean. Efficient estimation of word representations in vector space. arXiv preprint arXiv:1301.3781, 2013.

Zhen Wang, Jianwen Zhang, Jianlin Feng, and Zheng Chen. Knowledge graph embedding by translating on hyperplanes. In Proceedings of the AAAI Conference on Artificial Intelligence, volume 28 Issue 1, 2014.

Théo Trouillon, Johannes Welbl, Sebastian Riedel, Éric Gaussier, and Guillaume Bouchard. Complex embeddings for simple link prediction. In International conference on machine learning, pages 2071–2080. PMLR, 2016.

Tim Dettmers, Pasquale Minervini, Pontus Stenetorp, and Sebastian Riedel. Convolutional 2d knowledge graph embeddings. In Proceedings of the AAAI Conference on Artificial Intelligence, volume 32 Issue 1, 2018.

Komal Teru, Etienne Denis, and Will Hamilton. Inductive relation prediction by subgraph reasoning. In International Conference on Machine Learning, pages 9448–9457. PMLR, 2020.

Dobrik Georgiev, Marc Brockschmidt, and Miltiadis Allamanis. HEAT: hyperedge attention networks. CoRR, abs/2201.12113, 2022. URL https://arxiv.org/abs/2201.12113.

Mohammadamin Tavakoli, Alexander Shmakov, Francesco Ceccarelli, and Pierre Baldi. Rxn hypergraph: a hypergraph attention model for chemical reaction representation. arXiv preprint arXiv:2201.01196, 2022.

Romain Lopez, Jeffrey Regier, Michael B Cole, Michael I Jordan, and Nir Yosef. Deep generative modeling for single-cell transcriptomics. Nature methods, 15(12):1053–1058, 2018.

GTEx Consortium. The genotype-tissue expression (gtex) pilot analysis: multitissue gene regulation in humans. Science, 348(6235):648–660, 2015.

Alexander Lachmann, Zhuorui Xie, and Avi Ma’ayan. blitzgsea: efficient computation of gene set enrichment analysis through gamma distribution approximation. Bioinformatics, 38(8):2356–2357, 2022.

Mark D Robinson and Alicia Oshlack. A scaling normalization method for differential expression analysis of rna-seq data. Genome biology, 11(3):1–9, 2010.

Guido Van Rossum and Fred L Drake Jr. Python reference manual. Centrum voor Wiskunde en Informatica Amsterdam, 1995.

Adam Paszke, Sam Gross, Francisco Massa, Adam Lerer, James Bradbury, Gregory Chanan, Trevor Killeen, Zeming Lin, Natalia Gimelshein, Luca Antiga, Alban Desmaison, Andreas Kopf, Edward Yang, Zachary DeVito, Martin Raison, Alykhan Tejani, Sasank Chilamkurthy, Benoit Steiner, Lu Fang, Junjie Bai, and Soumith Chintala. Pytorch: An imperative style, high-performance deep learning library. In H. Wallach, H. Larochelle, A. Beygelzimer, F. d’Alché-Buc, E. Fox, and R. Garnett, editors, Advances in Neural Information Processing Systems 32, pages 8024–8035. Curran Associates, Inc., 2019. URL http://papers.neurips.cc/paper/9015-pytorch-an-imperative-style-high-performance-deep-learning-library.pdf.

Lukas Biewald. Experiment tracking with weights and biases, 2020. URL https://www.wandb.com/. Software available from wandb.com.

Charles R. Harris, K. Jarrod Millman, Stéfan J. van der Walt, Ralf Gommers, Pauli Virtanen, David Cournapeau, Eric Wieser, Julian Taylor, Sebastian Berg, Nathaniel J. Smith, Robert Kern, Matti Picus, Stephan Hoyer, Marten H. van Kerkwijk, Matthew Brett, Allan Haldane, Jaime Fernández del Río, Mark Wiebe, Pearu Peterson, Pierre Gérard-Marchant, Kevin Sheppard, Tyler Reddy, Warren Weckesser, Hameer Abbasi, Christoph Gohlke, and Travis E. Oliphant. Array programming with NumPy. Nature, 585(7825):357–362, September 2020. doi:10.1038/s41586-020-2649-2. URL https://doi.org/10.1038/s41586-020-2649-2.

F. Pedregosa, G. Varoquaux, A. Gramfort, V. Michel, B. Thirion, O. Grisel, M. Blondel, P. Prettenhofer, R. Weiss, V. Dubourg, J. Vanderplas, A. Passos, D. Cournapeau, M. Brucher, M. Perrot, and E. Duchesnay. Scikit-learn: Machine learning in Python. Journal of Machine Learning Research, 12:2825–2830, 2011.

Wes McKinney. Data Structures for Statistical Computing in Python. In Stéfan van der Walt and Jarrod Millman, editors, Proceedings of the 9th Python in Science Conference, pages 56–61, 2010. doi:10.25080/Majora-92bf1922-00a.

J. D. Hunter. Matplotlib: A 2d graphics environment. Computing in Science & Engineering, 9(3):90–95, 2007. doi:10.1109/MCSE.2007.55.

Michael L. Waskom. seaborn: statistical data visualization. Journal of Open Source Software, 6(60):3021, 2021. doi:10.21105/joss.03021. URL https://doi.org/10.21105/joss.03021.

Ilyes Khemakhem, Diederik Kingma, Ricardo Monti, and Aapo Hyvarinen. Variational autoencoders and nonlinear ica: A unifying framework. In International Conference on Artificial Intelligence and Statistics, pages 2207–2217. PMLR, 2020.

